# Detection, isolation and characterisation of phage-host complexes using BONCAT and click chemistry

**DOI:** 10.1101/2024.02.13.580147

**Authors:** P. Hellwig, A. Dittrich, R. Heyer, U. Reichl, D. Benndorf

## Abstract

Phages are viruses that infect prokaryotes and can shape microbial communities by lysis, thus offering applications in various fields. However, challenges exist in sampling, isolation, and predicting host specificity of phages. A new workflow using biorthogonal non-canonical amino acid tagging (BONCAT) and click chemistry (CC) allows combined analysis of phages and their hosts.

Replication of phage λ in *Escherichia coli* was selected as a model for workflow development. Specific labelling of phage λ proteins with the non-canonical amino acid 4-azido-L-homoalanine (AHA) during infection of *E. coli* was confirmed by LC-MS/MS. Subsequent tagging of AHA with fluorescent dyes via CC allowed the visualization of phages adsorbed to the cell surface by fluorescence microscopy. Flow cytometry enabled the automated detection of these fluorescent phage-host complexes. AHA-labeled phages were tagged with biotin for purification by affinity chromatography. The biotinylated phages could be purified and were infectious despite biotinylation after purification. Applying this assay approach to environmental samples would enable host screening without cultivation.

A flexible and powerful workflow was established to detect and enrich phages and their hosts. In the future, fluorescence-activated cell sorting or biotin purification could be used to isolate phage-host complexes in microbial communities.

## Introduction

Phages are viruses that infect prokaryotes and play a major role in the composition and evolution of microbial communities (Anderson, Brazelton and Baross, 2011; Heyer, Schallert and Siewert *et al*., 2019; Howard-Varona *et al*., 2017; Kristensen *et al*., 2010; Marsh and Wellington, 1994; Suttle, 2007). Phages are also considered a potential alternative to antibiotics for infection control (Clark and March, 2006; Fernández *et al*., 2021; Kutateladze and Adamia, 2010; Sieiro *et al*., 2020). Therefore, the identification and characterization of phages in natural, biotechnological and clinical areas, including patients, is of considerable interest.

DNA sequencing and genome annotation are crucial for phage detection and characterisation. However, associating an annotated phage sequence with a corresponding host is challenging and often impossible. Furthermore, DNA sequencing does not cover RNA phages (Fernández *et al*., 2021; Gregory *et al*., 2019; Paez-Espino *et al*., 2016) and cannot distinguish whether a phage genome is expressed or only integrated into the genome of the host cell. Heyer et al. (2019) demonstrated that metaproteomics could help to close these knowledge gaps by analysing the expression of phage proteins in microbial communities (Heyer, Schallert and Siewert *et al*., 2019). Nevertheless, analysing samples from complex microbial communities containing phages remains challenging due to their low contribution to the total biomass, the elaborate methods required for phage enrichment, and the lack of approaches to co-enrich corresponding host cells.

The enrichment and purification of low-abundant phage proteins and the visualisation of phage-host complexes are key to overcome these limitations. For example, Ohno *et al*. (2012) labelled phage DNA with 5-ethynyl-2′-deoxyuridine (EdU) and coupled fluorescent dyes to the labelled phage DNA. This approach was applied for the identification of the host specificity of phages. However, the use of this method is limited to DNA phages.

Furthermore, strong EdU labelling reduced infectivity, limiting the significance of plaque assays. Furthermore, the method required the injection of fluorescent phage DNA into the cell before detection by fluorescence microscopy or fluorescence-activated cell sorting (FACS). A more general approach, which would also cover RNA phages, is the labelling of phage proteins. In addition, phages could already be detected after adsorption to the surface of host cells. Hatzenpichler *et al*. (2014) showed the labelling of newly synthesised proteins in microbial communities using biorthogonal non-canonical amino acid tagging (BONCAT, Dieterich *et al*. (2006)) and click chemistry (CC, Kolb, Finn and Sharpless (2001). Pasulka *et al*. (2018) applied this approach to quantify the replication of phages by fluorescence microscopy. This approach was however not yet used to isolate phage-host complexes and purify phages from environmental samples.

As a remedy, we here present a workflow (Figure) to enrich and analyse phage-host complexes. The BONCAT workflow depends on incorporating 4-azido-L-homoalanine (AHA) into newly synthesised phage proteins. Combined with copper-free CC, fluorescent dyes or biotin can be attached to AHA-labelled phages without denaturation. Combining BONCAT and CC allowed the detection of labelled phage proteins adsorbed to the host surface and the specific enrichment of labelled phages. The workflow was evaluated using *E. coli* and phage λ as a well-established model system.

## Material and Methods

The workflow established is shown in Figure 1. A detailed description of the methods, including a step-by-step standard operation procedure, can be found in Supplementary Note 2: Material and Methods. Briefly, replication of phage λ was induced with mitomycin C (MMC) and the newly synthesised phage λ proteins were labelled with AHA (Figure 1, step 1 and 2). After centrifugation and filtration of the AHA-labelled phages, CC was used to attach fluorescent dyes or biotin (Figure 1, step 3a and b). Coupling of fluorescence dyes to phages enabled the detection of phages adsorbed to their host cells by fluorescence microscopy and FACS (in this work only flow cytometry) (Figure 1, step 4a). In addition, biotin coupling allowed the purification and enrichment of phages via affinity chromatography (Figure 1, step 4b). This enrichment allows the detection of the phages by liquid chromatography-mass spectrometry/mass spectrometry (LC-MS/MS). DNA/RNA sequencing would also be possible (the latter was not used here).

**Figure 1:**
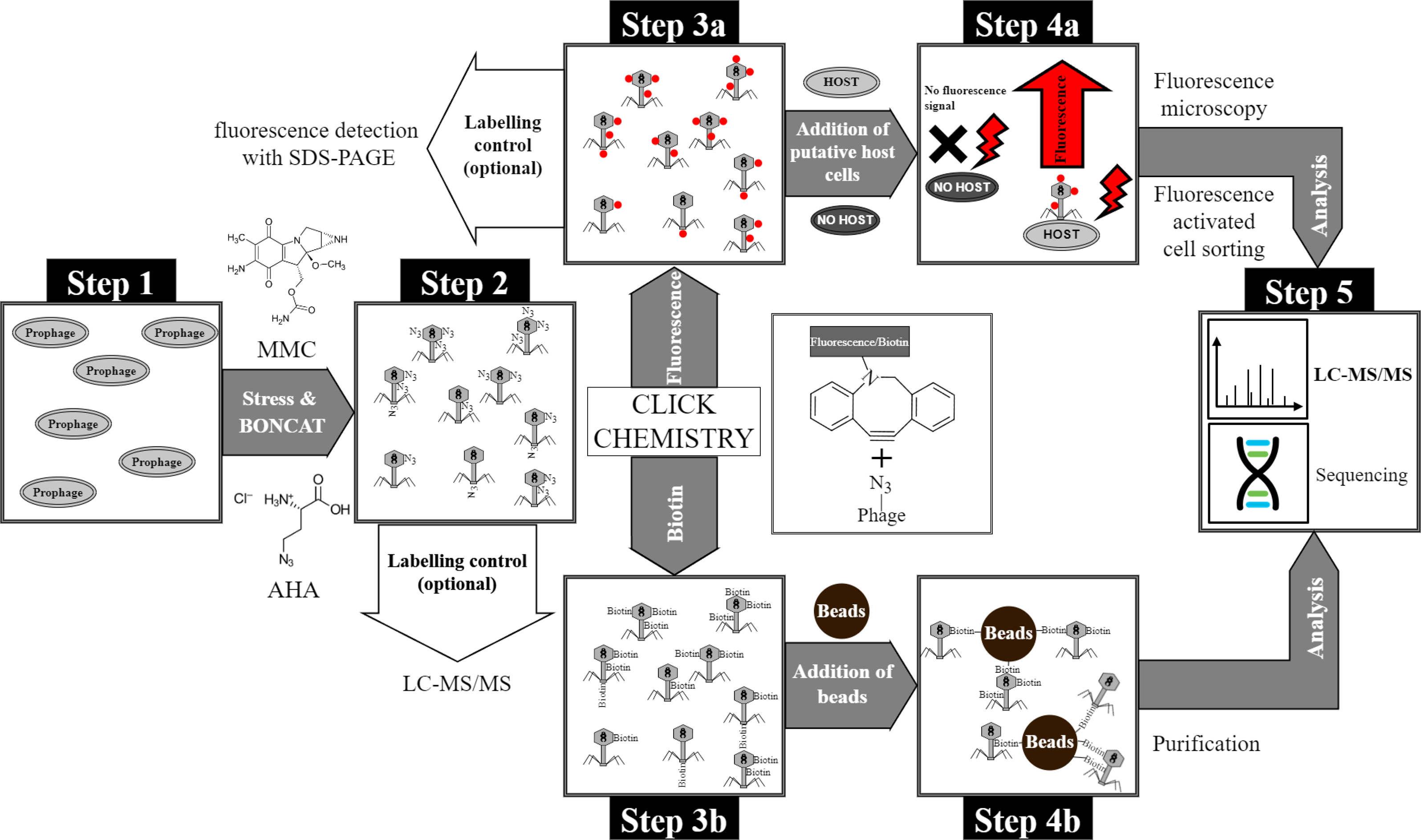
Overview of the BONCAT workflow for detection of phages and purification of phage-host complexes. **Step 1:** Cultivation of *E. coli* with genome-integrated phage λ, **Step 2:** Induction of phage replication with MMC and with subsequent incorporation of the non-canonical amino acid AHA into phage proteins. The incorporation of AHA into newly synthesised proteins was subsequently verified by LC-MS/MS. **Step 3:** Tagging of AHA-labelled phages with (**a**) fluorophores or (**b**) biotin using CC. In-gel detection is possible using a fluorescent dye (after CC) as quality control for CC. **Step 4a:** Incubation of fluorescent phages with putative host cells and identification/sorting of adsorbed phages by fluorescence microscopy and flow cytometry, respectively. **Step 4b:** Purification of biotin-labelled phages using magnetic beads (monomeric avidin beads). **Step 5:** Analysis of purified phages or phage-host complexes by LC-MS/MS or sequencing (not used in this study).

### Induction and replication of AHA-labelled phages

*E. coli* K12 (DSM 5911) with genome-integrated phage λ and *E. coli* K12 (DSM 5911) without phage integration were cultured in M9 minimal medium at 37 °C and 130 rpm overnight. Overnight cultures diluted to an optical density of 600 nm (OD_600_) ≈ 0.145 were used as inoculum to start new batches. After incubation of the bacteria for 1 h at 37 °C and 130 rpm, 0.5 µg/mL MMC and/or 0.1 mM AHA were added as indicated in Figure 2 and Supplementary Note 2: Table M1. Samples were taken to monitor bacterial growth, and OD_600_ was measured every hour. After 4 h or 6 h, phage λs were harvested by centrifugation of the culture (3000×g, 12 min, 4 °C). The supernatant was collected, and samples were adjusted to pH 7 with 1 M NaOH. Next, samples were clarified using a syringe with a filter cascade (5 µm, 1.2 µm, 0.8 µm, 0.45 µM, Sartorius AG). The cell pellets were not further analysed. For details, see Supplementary Note 2: Protocol S1.

**Figure 2:**
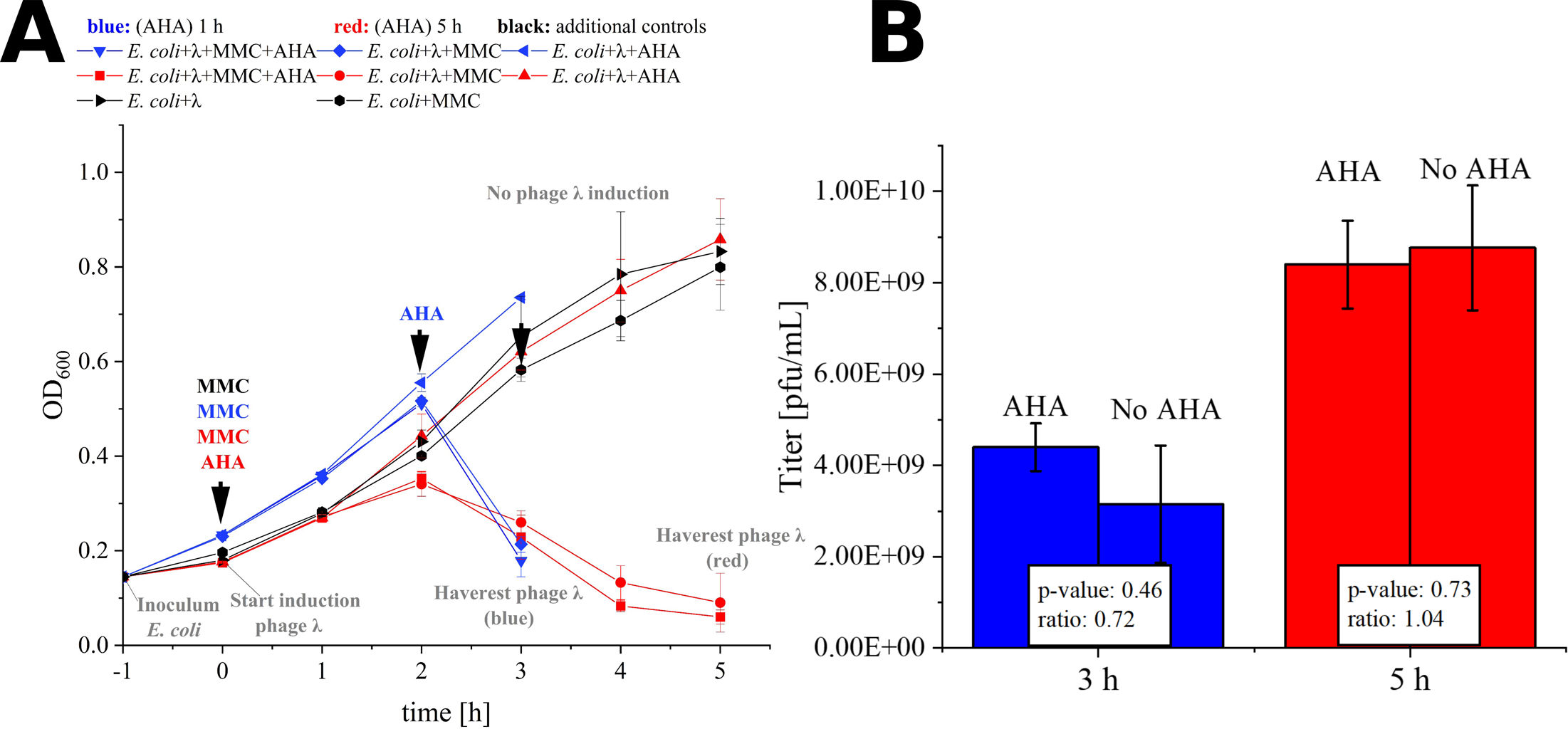
MMC induction and AHA labelling of phage λ. **A**) Time course of OD_600_ for 3 h resp 5 h phage λ induction. 4-azido-L-homoalanine (AHA) was added at the same time as MMC (AHA 5 h, red) or 1 h before the drop of the OD_600_ (AHA 1 h, blue). *E. coli* + λ: *E. coli* with integrated phage λ, see Supplementary Note 2: Protocol S1; **B)** Phage titer (pfu/mL) determined by plaque assay of the harvested phages from A. Mean and standard deviation of the phage λ titer for three independent experiments; differences in titer not significant (p>0.05; t-test with Benjamini-Hochberg correction). Raw data: see Supplementary Table 1.

### Plaque-assay

The phage suspensions were sequentially diluted (10^−6^ to 10^−7^), and 100 µL of each dilution was mixed with 4 mL of melted overlay agar and 100 µL of a fresh overnight culture of *E. coli* K12. Mixtures were poured onto a plate with underlay agar. Hardened agar plates were incubated at 37 °C overnight. Based on the number of plaques on plates, the phage titer was calculated: Number of plaques × 10 × reciprocal of dilution = pfu/mL. For details, see Supplementary Note 2: Protocol S2.

### Protein extraction, protein quantification, and sample preparation for LC-MS/MS

Proteins from 4 mL of each phage suspension were extracted with chloroform-methanol. Then, proteins were resuspended in 1 mL 8 M urea buffer. The protein concentration was quantified using amido black assay. 25 µg of total protein was used for Filter Aided Sample Preparation (FASP) digestion with MS-approved trypsin (1:100 µg protein) (Heyer, Schallert and Büdel *et al*., 2019). The resulting peptide solutions were dried in a vacuum centrifuge and solubilised in 75 µL loading buffer A (LC-MS water and 0.1% trifluoroacetic acid (TFA)). For details, see Supplementary Note 2: Protocol S3.

### LC-MS/MS

LC-MS/MS analysis was performed using an UltiMate® 3000 nano splitless reversed-phase nanoHPLC (Thermo Fisher Scientific, Dreieich) coupled online to a timsTOF™ pro mass spectrometer (Bruker Daltonik GmbH, Bremen). For details, see Supplementary Note 2: Protocol S3.

### Identification of AHA incorporation using MASCOT

MS/MS raw data files were processed with the Compass DataAnalysis software (version 5.3.0, Bruker Corporation, Bremen, Germany) and converted to Mascot Generic Files (.mgf). The files were uploaded to MASCOT Daemon (Version 2.6.0) (Perkins *et al*., 1999) and searched against a filtered UniProt database containing only *E. coli* K12 (taxonomy_id: 83333, 23.03.2023) and “Bacteriophage lambda” (taxonomy_id: 10710, 23.03.2023) entities. The following modifications were used: oxidation of methionine, carbamidomethyl, AHA, and reduced AHA (see Supplementary Note 2: Table M2).

### Fluorophore/biotin tagging of BONCAT phages by click chemistry

Phage suspensions were collected on a 100 kDa filter via centrifugation (5 min, 3,500×g, RT). Afterwards, phosphate-buffered saline (PBS) was added and samples were centrifuged again (5 min, 3,500×g, RT). 100 mM iodoacetamide in PBS was added, and samples were incubated in the dark at 37 °C for 1 h. Afterwards, 0.15 µM dibenzylcyclooctyne (DBCO)-cyanin 5.5 or 0.15 µM DBCO-Alexafluor 555 or 0.15 mM DBCO-PEG_4_-biotin were added. Next, samples were incubated in the dark for 30 min at 37 °C, washed thrice with PBS, and resuspended in 1 mL PBS. Finally, phage suspensions were transferred to 1.5 mL LoBind^®^ tubes and stored at 4 °C in the dark. For details, see Supplementary Note 2: Protocol S4.

### Specific adsorption of labelled phages to host cells

*E. coli* and *Pseudomonas fluorescens* (DSM 50090) were cultured in standard nutrient broth (plus 5 mM MgSO_4_) at 37 °C, 130 rpm overnight. Next, bacteria were diluted with fresh standard nutrient broth in sterile 1.5 mL tubes to 1.80 × 10^7^ cells/mL and incubated for 20 min at 30 °C and 600 rpm. Fluorescent phages were added to adjust a multicity of infection (moi) ≈ 2. As a control, only medium was added to the bacteria. 200 µL samples were taken after 0 min, 10 min, 20 min, 30 min, and 60 min. The samples were immediately centrifuged (5 min, 16,400×g, 4 °C). The supernatants of the samples were removed, and cell pellets were immediately fixed with 4% formaldehyde in PBS for 1 h at 4 °C. The fixation solution was removed by centrifugation (5 min, 16,400×g, 4 °C). Cells were resuspended in PBS and stored at 4 °C. For details, see Supplementary Note 2: Protocol S4.

### Fluorescence microscopy

Phage-host complexes were visualised with an Imager.M1 fluorescence microscope (Carl Zeiss, Jena, Germany) using a 100X objective (EC-Neoflur 100x/1.3 Oil Ph3) and phase contrast. For details, see Supplementary Note 2: Protocol S4.

### Flow cytometry

Flow cytometric analysis was performed using a FACS Canto II equipped with three lasers (405 nm, 488 nm, 663 nm), Firmware Version 1.47 (BD Biosciences, Franklin Lakes, NJ, USA). The data were analysed with the software FlowJo™ (BD Biosciences, 10.8.1). For details, see Supplementary Note 2: Protocol S4.

### Native purification of biotinylated phages via magnetic beads

Biotinylated phages were purified with BcMag™ Monomeric Avidin Magnetic Beads (Bioclone, MMI-101) kit according to the manufacturer’s instructions. After binding of the phages, beads were washed with PBS. The supernatant of each washing step was collected for further analysis (fraction “Washing phase”). The biotinylated phages bound to the beads were eluted with 2 mM biotin and collected in a new tube (fraction “Elution”). Lastly, the beads were boiled at 60 °C for 5 min with an SDS-buffer, and the supernatant was collected for further analysis (fraction “SDS-boiled”). All fractions were analysed with an untreated control (fraction “not purified”) with SDS-PAGE. The phage titer in the ‘Elution’ was also determined with plaque assay. For details, see Supplementary Note 2: Protocol S4.

### SDS-PAGE

SDS-PAGE was performed with 1 mm SDS-PAGE gels with 12% separation and 4% stacking gel (Laemmli, 1970). For details, see Supplementary Note 2: Protocol S4.

### Staining and scanning of SDS gels loaded with biotinylated or fluorescent proteins

After electrophoresis and fixation gels with fluorescent proteins were scanned with Licor Odyssey ODY-2600 (LI-COR Biosciences - GmbH) or Typhoon Trio Variable Mode Imager System (GE Healthcare). Subsequently, the gels were counterstained with Coomassie staining solution overnight and scanned with a Biostep ViewPix900 scanner (Seiko Epson Corporation) (Supplementary Note 2: Table M 3). Gels with biotinylated proteins were fixed, stained with Coomassie, and scanned with a Biostep ViewPix900 scanner (Seiko Epson Corporation) (Supplementary Note 2: Table M 3).

### In-gel digestion

The method was performed as described in Heyer, Schallert and Büdel *et al*., 2019. For each protein band isolated from an SDS gel, 1 µg of protein content was assumed to calculate the amount of MS-approved trypsin (1 µg trypsin: 100 µg protein).

### Replicates, biostatistics, and visualisation

All experiments were performed in biological triplicates or as indicated. R-Statistics (version 4.1.2) with R studio (version 2021.09.1 Build 372) was used for statistical analysis. Normal distribution was confirmed by the Shapiro-Wilk test; for group-wise differences, a t-test with Benjamini-Hochberg correction was used.

## Results and Discussion

### BONCAT labelling of phage λ in *E. coli*

Efficient phage replication induced by MMC was a critical precondition for the subsequent labelling of phages with AHA. The addition of MMC to exponentially growing *E. coli* resulted in a growth arrest at 2 h and a subsequent decrease of biomass (OD_600_), indicating cell lysis and phage replication (Figure 2 A). Based on OD_600_, the addition of AHA for labelling did not reduce phage replication. Similar phage titers confirmed this result for incubations with and without additions of AHA (Figure 2 B, p>0.05). Earlier haverest at 3 h resulted in lower phage titers, showing that phage replication was still ongoing until final sampling at 5 h.

In summary, AHA addition did neither inhibit the MMC-induced production of phage λ in *E. coli* nor the production and infectivity of phage λ. This is consistent with recent studies investigating the impact of AHA addition on the growth of *E. coli* (Landor *et al*., 2022; Steward *et al*., 2020).

### Verification of the incorporation of AHA by LC-MS/MS

The successful labelling of proteins with AHA was subsequently confirmed using LC-MS/MS. Here, the incorporation of AHA instead of methionine caused specific mass shifts of tryptic peptides. Overall, LC-MS/MS allowed the assignment of 5,046 ± 1,200 peptide spectrum matches (PSMs) related to phage λ proteins (Figure 3 A). After 5 h labelling with AHA, 271 ± 50 PSMs showed incorporation of AHA instead of methionine. This accounted for 5.68% ± 0.23% AHA labelled PSMs compared to all PSMs detected for phage λ. Since AHA is only incorporated instead of methionine, the calculation of the incorporation of AHA in all methionine-containing PSMs (51.82% ± 1.15% of all PSMs) should be considered as reference. Taken this into account, the degree of labelling with AHA is doubled. Interestingly, shorter labelling with AHA (1 h) resulted in a similar incorporation of AHA (5.24% ± 2.14% of all PSMs) (Figure 3 A, Supplementary Table 2), showing that AHA incorporation started soon after addition. Compared to eukaryotic cells, where AHA is only incorporated at 1 out of 400-500 methionine sites (Calve *et al*., 2016; Kiick *et al*., 2002; Ngo *et al*., 2009; van Bergen, Heck and Baggelaar, 2022), the incorporation of AHA observed for phage λ was higher. This has several advantages. (i) It provides many reactive sites for subsequent coupling of fluorophores or affinity tags. It may partially compensate (ii) for the lower occurrence of methionine in some other phages or (iii) for the lower incorporation rate of AHA in other bacterial species. In the latter case, the AHA concentration in the supernatant could be further increased (up to 1 mM) or AHA could be added continuously at low concentrations to reduce its impact on the physiology of host cells (Hatzenpichler *et al*., 2014; Landor *et al*., 2022; Steward *et al*., 2020).

**Figure 3:**
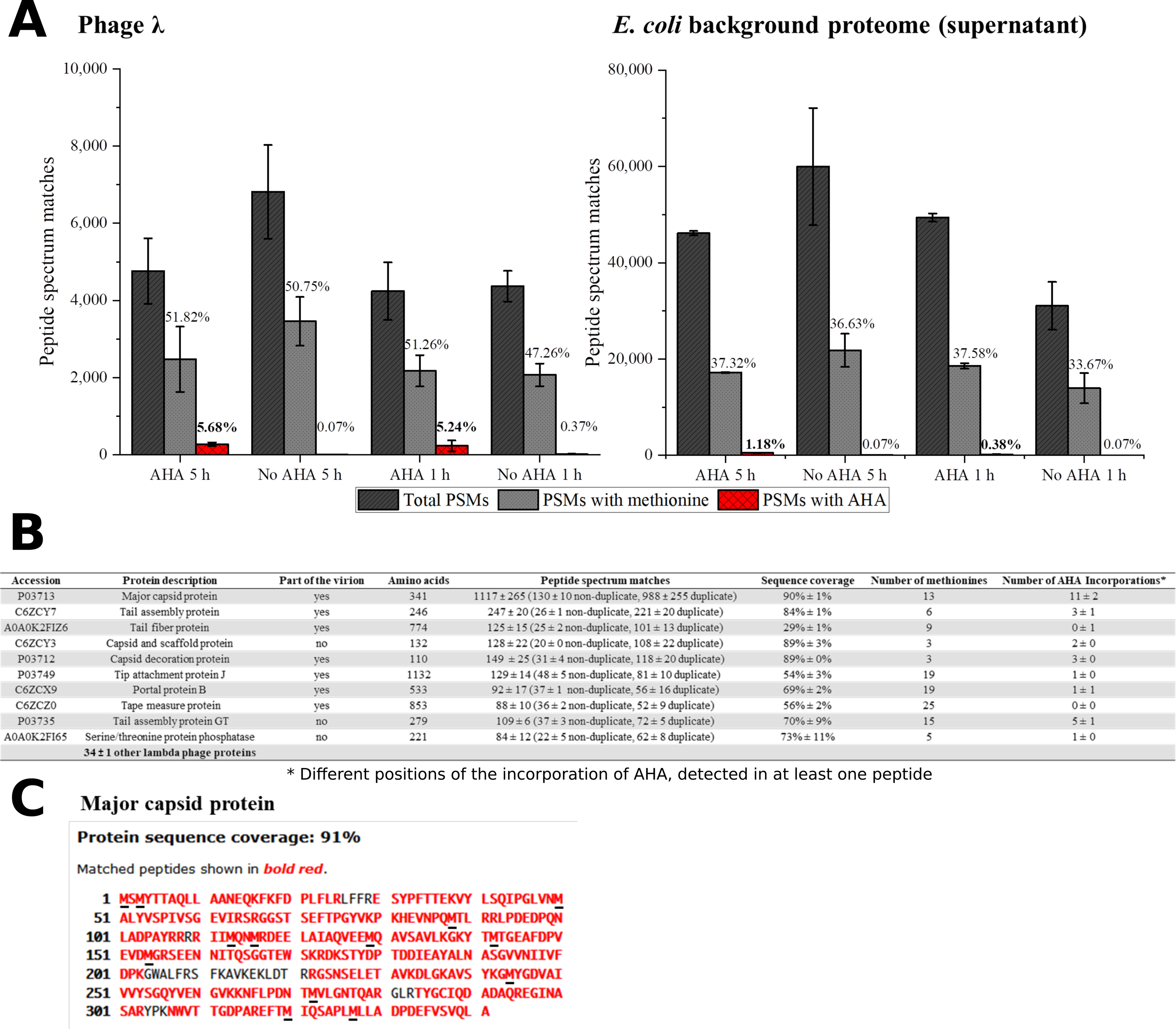
Screening for AHA containing PSMs from phage λ and the *E. coli* background proteome. PSMs were identified by data-dependent acquisition LC-MS/MS using the MASCOT search engine. **A)** A total number of phage λ and *E. coli* (not cell pellet); only PSMs of the same sample were analysed. AHA 5 h, AHA 1 h: AHA incubation times of 1 h and 5 h; No AHA 5 h, No AHA 1 h: controls. PSMs with methionine indicate the percentage of methionine-containing PSMs of all identified PSMs. PSMs with AHA indicate the percentage of AHA-containing PSMs of all PSMs. Mean and standard deviation of three independent experiments (Supplementary Table 2). **B)** The top 10 identified phage λ proteins containing AHA after MASCOT search for sample AHA 5 h (3 replicates) were analysed—the order of the top 10 phage λ proteins corresponds to replicate 1. PSM: Mean and standard deviation of three replicates. **C)** The major capsid protein was the most abundant protein detected in all samples (90 %±1 sequence coverage). Black underlined letters in the amino acid sequence shows the position of methionines. The possibility of AHA incorporation into the major capsid protein was confirmed for all positions of methionine of replicate 1.

Detailed analysis of PSMs allowed to identify 44 ± 1 different phage λ proteins (Figure 3 B) associated with the infection cycle. The most abundant phage protein was the major capsid protein, where AHA was incorporated in all methionine positions (Figure 3 C). The objective of incorporating AHA should be to provide adequate binding sites for the CC while retaining the functionality of phages. The incorporation of AHA failed only in 1 out of the top 10 identified phage λ proteins (Figure 3 B). This result could indicate that the incorporation of AHA impacts the stability or the function of labelled proteins, potentially interfering with the infectivity of the phages (Landor *et al*., 2022). However, according to the results obtained from plaque assays, labelling with AHA at the given concentration does not significantly affect titers (Figure 2 B). Heterogeneity of incorporation of AHA in other proteins of phage λ might be caused either by selective incorporation of AHA or by removal of dysfunctional/misfolded proteins after protein synthesis.

During phage-induced cell lysis, the phage harvest may become contaminated with *E. coli* proteins, which could interfere with the subsequent dye or biotin labeling steps of the CC. LC-MS/MS analysis of phage harvest after purification by centrifugation thus showed the presence of a relatively large number (46,636 ± 11,934 PSMs) of *E. coli* background proteins containing 1.18% ± 0.10% AHA labelled PSMs for 5 h AHA labelling, and 0.38% ± 0.15% AHA labelled PSMs for 1 h AHA labelling. Therefore, shorter labelling with AHA (1 h) in a later phase of infection (2 h after the addition of MMC) should be preferred due to undesired labelling of *E. coli* background proteins (Figure 3 A) despite lower labelling efficiency. However, interfering background proteins could also be removed by CsCl centrifugation or PEG precipitation (Boulanger, 2009; Nasukawa *et al*., 2017; Yamamoto *et al*., 1970). Nevertheless, every additional purification step might also reduce the yield of phages (Carroll-Portillo *et al*., 2021).

In summary, both tested AHA incubation periods allowed the successful AHA-labelling of the phages. However, the parallel incubation of cells with the phage replicating inducer (here MMC) and AHA is more practical, especially for cultures with unknown cell growth dynamics and phage replication kinetics. Therefore, the 5 h incubation period with simultaneous MMC and AHA addition was used in the further course of this study.

### Fluorescence tagging of AHA-labelled phages

AHA-labelling was a precondition for the attachment of fluorescent dyes by CC to identify newly synthesised proteins by SDS-PAGE and fluorescence microscopy (Figure 1 Step 3a) (Dieterich *et al*., 2007; Hatzenpichler *et al*., 2014; Pasulka *et al*., 2018).

Previously published protocols for CC apply precipitation with ethanol to remove excess reagents. However, the denaturation of phages by ethanol precipitation should be omitted for subsequent fluorescence microscopy. Therefore, the protocol to tag the AHA-labelled phages with fluorophores was adapted, and all CC and washing steps were performed using a 100 kDa filter, retaining the native phages in the supernatant (Bichet, Patwa and Barr, 2021; Bonilla *et al*., 2016; Erickson, 2009; Hietala *et al*., 2019). Ultracentrifugation was not considered here, as pelleting phages was considered too time-consuming and potentially reducing overall yield. The newly established filter-based protocol for tagging phages with DBCO Alexafluor (AF) 555 allowed the successful detection of fluorescence in SDS-PAGE (see Supplementary Note 1: Figure S 1). Two fluorescent bands with molecular weights of approximately 37 kDa and 60 kDa were detected for both labelling conditions (1 h and 5 h labelling with AHA). In contrast, the control showed no fluorescent bands. The 37 kDa band could correspond to the highly labelled major capsid protein, and the 60 kDa band to the portal protein B. Further, these fluorophore-tagged phages are termed “AF555 phages”. Alternatively, AHA-labelled phages were coupled to DBCO Cyanin (CY) 5.5 by CC (see Supplementary Note 1: Figure S 4). The fluorescence gels showed higher fluorescence intensity with additional bands besides the two main bands at 37 kDa and 60 kDa. In the following, these fluorophore-tagged phages are termed “CY5.5 phages”.

In summary, the phages had sufficient binding sites for detectable fluorescence tagging via CC. This also confirms the results of the MS measurements, where a high level of incorporation with AHA was found. In addition, sufficient phages could be recovered from the filters for phage protein detection via Coomassie stain and fluorescence. This should also allow fluorescence microscopy detection, which will be verified below.

### Detection of phage-host complexes via fluorescence microscopy

Next, the binding of AF555 phages to *E. coli* was approved by fluorescence microscopy. AF555 phages were incubated for 30 min with *E. coli* or *P. fluorescens* as negative control.

Neither *P. fluorescens* nor *E. coli* showed background fluorescence at the selected wavelength (Figure 4 A). *E. coli* incubated with AF555 phages emitted fluorescence (Figure 4 A), whereas *P. fluorescens* incubated with AF555 phages in most cases did not emit fluorescence. Despite their small size, AF555 phages were even visible as red dots on the surface of *E. coli* cells (Figure 4 B). This even applied to fluorescent phages in the supernatant (Supplementary Note 1: Figure S 3, see Pasulka *et al*. (2018)).

**Figure 4:**
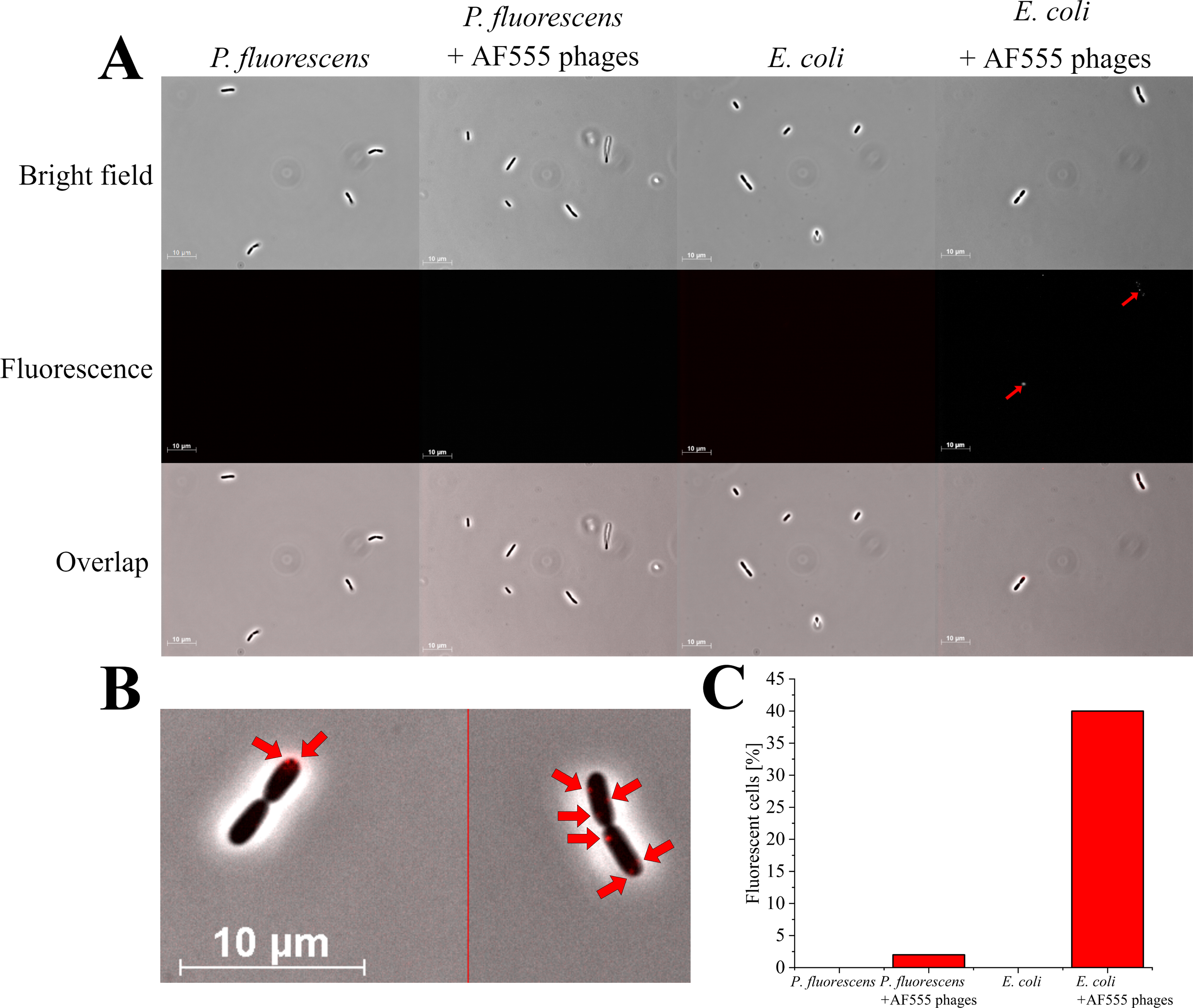
Fluorescence microscopy of *E. coli* and *P. fluorescens* (control) with AF555 phages. Cells were incubated for 30 min with and without AF555 phages. All pictures were taken with an Imager M1 fluorescence microscope (Carl Zeiss, Jena) using the software AxioVision (Version 4.8.2 SP3); brightfield and fluorescence (excitation 546/12 nm; beamsplitter: FT 560; emission 575-640 nm), 1000x, phase contrast. **A**) Representative pictures of *P. fluorescens* and *E. coli* cells with and without AF555 phages after incubation for 30 min; only *E. coli* cells show a positive fluorescence after the addition of AF555 phages (red arrow). **B)** *E. coli* cells with attached AF555 phages (red arrow, enlarged from overlap in A). **C)** Percentage of fluorescent cells after scanning in greyscale mode (see Supplementary Note 1: Figure S 2).

About 40% of *E. coli* cells showed AF555 phages-specific fluorescence signals after 30 min incubation, whereas only 2% of *P. fluorescens* showed a fluorescence signal (Figure 4 C). The small proportion of AF555 phages bound to *P. fluorescens* could be explained by the unspecific binding of phages to glycans of the extracellular membrane on many gram-negative bacteria (Dennehy and Abedon, 2021; Maffei *et al*., 2021) that are similar to carbohydrates of the *E. coli* membrane.

In summary, a workflow for AHA-labelling and CC-based addition of DBCO AF555 from phage λ was established. AF555 phages specifically bound to their host cells with little negative impact on infectious titer.

### Quantification of fluorescent phage-host complexes via flow cytometry

Fluorescence microscopy was used to monitor the absorption of AF555 phages on their host cells (Figure 4). In addition, flow cytometry was applied for high-throughput analysis to quantify phage-host interaction (Figure 1 Step 4a).

First, CY5.5 phages were incubated with *E. coli* or *P. fluorescens* for up to 60 min (Figure 5 A “λ”)*. E. coli* and *P. fluorescens* incubated without CY5.5 phages served as control (Figure 5 A “C”). Pure bacteria (without phages) and pure CY5.5 phages were used to define gates excluding clumped bacteria and unbound fluorescent phages from counting as positive signals for phage-host interaction analysis (see Supplementary Note 1: Figure S 5).

**Figure 5:**
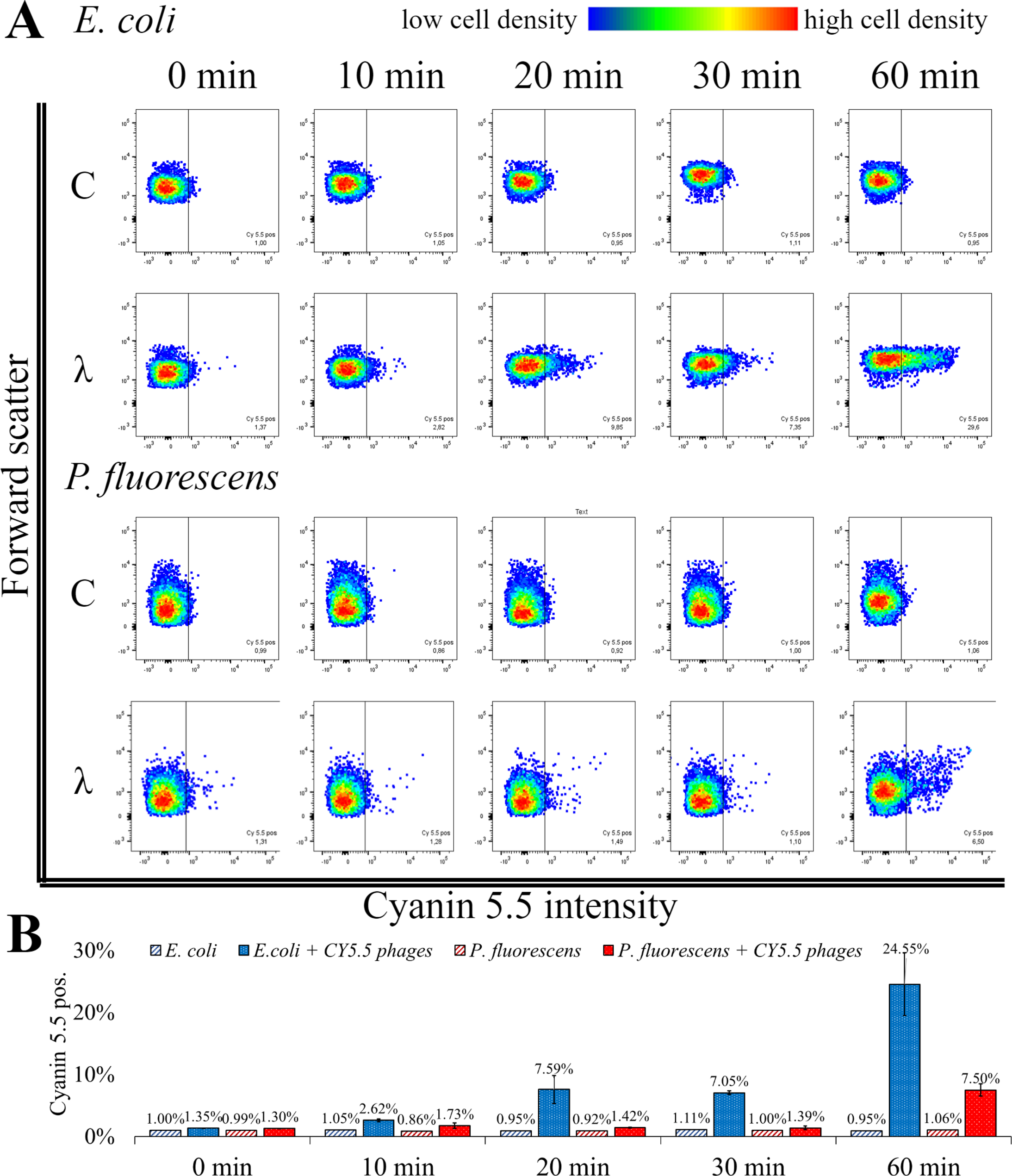
Flow cytometric analysis of phage-host complexes. **A)** *E. coli* or *P. fluorescens* were incubated with CY5.5 phages for the indicated period (λ); control (C): without the addition of CY5.5 phages. Bacteria were harvested and treated with 4% formaldehyde for fixation. The fluorescence intensity was determined for n = 10,000 bacteria per condition. Forward scatter as an indicator of cell size was plotted against the intensity of cyanin 5.5 fluorescence. The threshold for background fluorescence of bacteria incubated without phages was set to about 1% (vertical line in the scatter plots). **B)** Mean percentage of cyanin 5.5 positive phage-host complexes from two independent biological replicates with standard derivation. The third replicate showed different adoption kinetics and the raw data can be found in Supplementary Table 3: FACS analysis.

The percentage of fluorescent *E. coli* incubated with CY5.5 phages increased from 1.35% ± 0.04% to 24.55% ± 7.14%. A rapid increase in fluorescence of *E. coli* from 7.05% ± 0.42% to 24.55% ± 7.14% was observed between 30 min and 60 min incubation with CY5.5 phages. In contrast, the fluorescence of *P. fluorescens* did not increase significantly within the first 30 min of incubation using CY5.5 phages. After 60 min of incubation of *P. fluorescens* with CY5.5 phages, 7.50% ± 1.41% of the cells were fluorescent, indicating unspecific binding of CY5.5 phages due to the extended incubation time (Figure 5). Unspecific adsorption of CY5.5 phages to non-host cells such as *P. fluorescens* might cause false positive results. Therefore, short incubation times are suggested. Alternatively, unspecific binding could be minimised by extensive washing steps with PBS after harvest.

In our analysis of the three biological replicates, we observed a slower adsorption of CY5.5 phages in one replicate, where the increase in fluorescence from *E. coli* after CY5.5 phage addition increased from 1.12% to 4.13% after 60 min, while the fluorescence of *P. fluorescens* remained at 1.17% (Supplementary Table 3). A longer incubation time might have also increased the fluorescence signal in this experiment, similar to the other replicates. Therefore, we recommend always to perform replicates for the analysis.

In the case of low fluorescence signals, the number of phages bound to host cells could be increased by testing a higher moi. Since phage infections follow a Poisson distribution, incubation of bacteria with higher moi could lead to stronger fluorescence since more phages per cell surface are to be expected (Arkin, Ross and McAdams, 1998; Ellis and Delbrück, 1939; Kourilsky, 1973; Marcelli *et al*., 2020). In particular, for unknown phage titers, the optimal adsorption conditions must be determined by analysing different ratios of cells and phages.

Counterstaining of the cells with another fluorescent dye may also help when analysing more complex samples (Reichart *et al*., 2020).

In summary, the analysis of fluorescent phage-host complexes by flow cytometry confirmed the results obtained by fluorescence microscopy (Figure 4) with the advantage of high-throughput quantification. Furthermore, it offers options for the enrichment and isolation of fluorescent phage-host complexes by fluorescence-associated cell sorting (FACS) in follow-up studies.

### Purification of biotinylated phages via magnetic beads

CC of AHA-labelled proteins allows coupling of affinity tags, such as biotin, permitting specific enrichment of tagged proteins with corresponding binding partners, such as avidin (Wilchek and Bayer 1990). Enrichment of intact biotinylated phages from complex cultures would allow subsequent analyses of the isolated phages, including DNA/RNA sequencing, LC-MS/MS-based proteomics, or follow-up infection experiments (Figure 1 Step 3b and 4b).

AHA labelled phages were tagged with DBCO-PEG4-biotin (biotinylated phages) and purified with magnetic beads functionalised with monomeric avidin. The different fractions obtained were analysed by SDS-PAGE (Figure 6 A). The biotinylated phages eluted easily using a surplus of biotin (Figure 6 A “biotinylated phages” SDS gel lane “Elution”). In contrast, the elution of non-biotinylated phages (control) failed (Figure 6 A “non-biotinylated phages” SDS gel lane “Elution”). The low protein content of the collected washing fractions of beads (Figure 6 A SDS gel lanes “Washing phase”) indicated that too little protein is released by the mild washing of beads with PBS. In contrast, boiling the beads with an SDS-buffer after the elution step removed many proteins from the beads, indicating an unspecific binding to the bead surface, which is independent of the biotinylation (Figure 6 A SDS gel lanes “SDS-boiled”). However, the unspecific binding of proteins to the monomeric avidin beads does not seem to affect the purification of phages since exclusively biotinylated phages were collected after the addition of biotin (Figure 6 A “biotinylated phages” SDS gel lane “Elution”).

**Figure 6:**
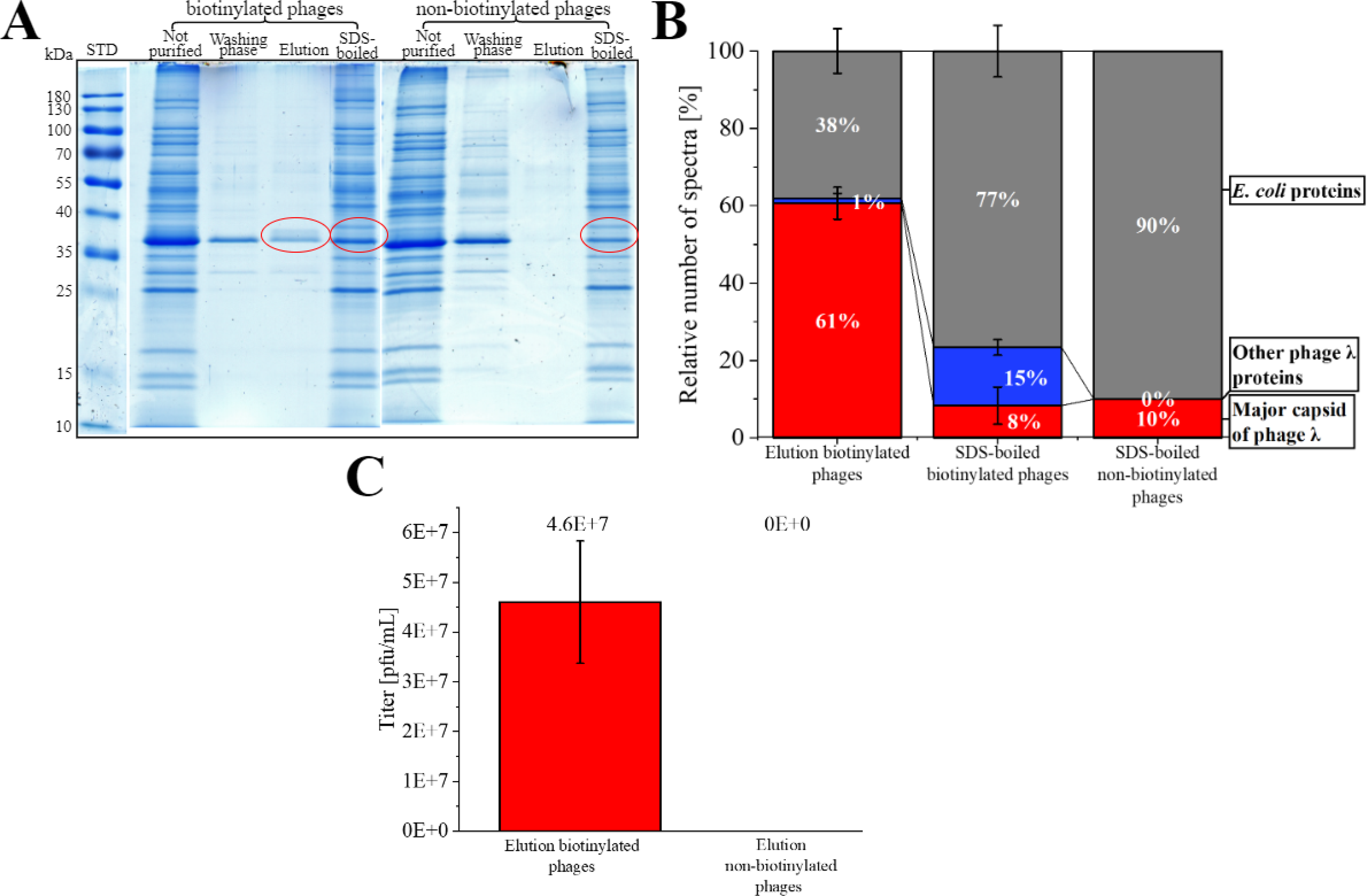
Purification of biotinylated phages with magnetic monomeric avidin beads. **A)** SDS gel from different fractions collected during purification of phages (biotinylated or non-biotinylated) stained with Coomassie blue. The fraction “not purified” corresponds to the control (sample before the addition of beads). The “washing phase” comprises fractions of all washing steps of the beads with PBS. “Elution” includes all proteins eluted from avidin beads with a surplus of biotin. The fraction “SDS-boiled” is the collected supernatant of avidin beads after boiling (5 min, 60 °C) with SDS buffer. Red circles indicate protein fractions that were further analysed by LC-MS/MS. For the original gels, see Supplementary Note 1 Figure S6. STD: protein standard (Thermo Scientific, PageRuler Prestained Protein Ladder #26616) **B)** Relative percentage of the spectra measured with LC-MS/MS after in-gel digestion of the protein fractions marked in **A**. MS data were screened for the major capsid protein from phage λ, other phage λ proteins, and *E. coli* proteins from the background of the phage λ suspension. For raw data, see Supplementary Table 4 MS in-gel. **C)** Plaque titer from the elution fraction of biotinylated and non-biotinylated phages; three independent experiments with mean and standard derivation below the detection limit for the elution of non-biotinylated phages.

The most abundant protein from the “Elution” of the biotinylated phages had a molecular weight of about 37 kDa. It corresponded to the major band of AF555 and CY5.5 phages identified by SDS-PAGE (Figure 6 A). LC-MS/MS confirmed that this band mostly contained the major capsid protein of phage λ (Figure 6 B red) demonstrating the specific elution of biotinylated phages from avidin beads. The additional analysis of the SDS boiled fractions showed a low proportion of major capsid protein but a high proportion of *E. coli* PSMs.

Obviously, destroying the beads with SDS mostly released background proteins (Figure 6 B), whereas a mild elution with 2 mM biotin is highly specific. Blocking the beads with amino acids or gelatin could be considered in future applications to prevent the unspecific binding of host proteins.

Next, plaque assays were performed to control the infectivity of the collected phages. Here, biotinylated phages eluted from beads showed infectivity with 4.60E+07 pfu/mL ± 1,23E+07 pfu/mL (Figure 6 C), whereas the elution fraction of beads loaded with unlabelled showed no plaques. The missing plaques in the unlabelled experiment confirmed the specific binding of biotinylated phages, as already concluded from LC-MS/MS data. In summary, it is possible to tag phages with biotin using CC without compromising their infectivity. Biotin tagging allows the purification of the biotinylated phages with monomeric avidin beads. The specific bead-bound phages could also be used (after optimisation) for specific host screening, where the phages bind the hosts, and the phage-host complexes are specifically released from the beads by an excess of biotin (mild condition).

### Future application of the established workflow in microbial ecology and personalised medicine

The new workflow established enabled fluorescence labelling of phage λ for subsequent monitoring by fluorescence microscopy and flow cytometry. A specific enrichment of infectious biotin-labelled phage λ fractions is possible using monomeric avidin beads. Further phage-host systems could be tested for future applications in microbial ecology and personalized medicine. As BONCAT approaches have been applied to a wide range of species (e.g., (Babin *et al*., 2017; Franco *et al*., 2018; Metcalfe *et al*., 2021; Pasulka *et al*., 2018)) no major difficulties in the transfer are anticipated. Fluorescent labelling of phages from phage collections would enable high throughput screening using flow cytometry for alternative hosts in bacterial strain collections or the personalised selection of phages for phage therapy using the pathogenic isolate from the patient as targets. The screening could also be widened to non-cultivable bacteria enriched from environmental samples. Flow cytometry-based cell sorting and subsequent sequencing or proteomics could support the identification and description of new hosts for phages already available in phage collections. BONCAT and induction of phage replication by MMC or other environmental stressors could also be applied to microbial communities (e.g. (Howard-Varona *et al*., 2017; Jiang and Paul, 1998; Rossi *et al*., 2022)). Phages could be separated from cells by filtration or ultracentrifugation for subsequent labelling with fluorescent dyes or affinity tags. Afterwards, flow cytometry and cell sorting can be applied to identify and characterise the corresponding hosts, including non-cultivable bacteria from the same microbial community. Alternatively, biotinylated phages previously immobilised on monomeric avidin magnetic beads can enrich corresponding hosts for subsequent sequencing and characterisation by adsortption on the surface of host cells. Although the application to microbiomes sounds very ambitious, it holds great potential for the identification and monitoring of phage-host interactions in their natural environment. In particular, binding conditions should be optimised for analysis of complex samples from the environment to reduce the risk of false-positive assignments.

### Conclusion

A workflow for the analysis of phage λ replication in *E. coli* and the detection and purification of fluorescent phage-host complexes was established. First, phages were labelled with AHA using BONCAT. Second, labelled phages were tagged with either fluorescent dyes or biotin using CC. Using BONCAT followed by CC, is a novel strategy and flexible tool for studying microbial communities (Hatzenpichler *et al*., 2020). The established method was exemplarily applied to pure cultures but can serve as a basis for analysing the function of phages in diverse, complex microbiological communities, including environmental or patient samples. Furthermore, fluorescent phages could be applied for specific screening of phage libraries for therapy of infectious diseases.

## Supporting information

Supplementary_note_2_Material_and_Methods_SOPs

Supplementary table 1 induction and plaque assay

Supplementary table 2 AHA incorporation LC-MS

Supplementary table 3 FACS analysis

Supplementary table 4 MS ingel

## Acknowledgment

We thank Prof. Dr. Andreas Kuhn (University of Hohenheim, Germany) for providing *E. coli* with integrated phage λ and the helpful feedback to the manuscript and Helga Tietgens (Max Planck Institute for Dynamics of Complex Technical Systems Magdeburg, Germany) for support in fluorescence microscopy.

## Supplementary

Supplementary Table 1 induction and plaque assay

Supplementary Table 2 aha incorporation LC-MS

Supplementary Table 3 FACS analysis

Supplementary Table 4 MS in gel

Supplementary Note 1 figures and tables

Supplementary Note 2 material & methods

## Abbreviations

AF: alexafluor
AHA: 4-azido-L-homoalanine
BONCAT: biorthogonal non-canonical amino acid tagging
CC: click chemistry
CID: collision induced dissociation
CY: cyanin
DBCO: dibenzylcyclooctyne
DDA: data dependent acquisition
DIA: data independent acquisition
EdU: 5-ethynyl-2′-deoxyuridine
FACS: fluorescence activated cell sorting
FASP: Filter Aided Sample Preparation
LC-MS/MS: liquid chromatography-mass spectrometry/mass spectrometry
MMC: mitomycin c
Moi: multiplicity of infection
ncAA: non-canonical amino acids
OD: optical density
PASEF: parallel Accumulation Serial Fragmentation
PBS: phosphate-buffered saline
PSM: peptide spectrum match
Rpm: rounds per minute
RT: room temperature
TIMS: trapped ion mobility spectrometry
TFA: trifluoroacetic acid

