## supplementary_note_1_figures_and_tables for "Detection, isolation and characterisation of phage-host complexes using BONCAT and click chemistry"

Table S 1: t-test of titer from AHA-labelled and non-labelled induced phage λs in *E. coli* with Mitomycin C. The t- test was performed with R studio (see Material and Methods). The p-value indicates no significant difference between the groups AHA R1-3 and No AHA R1-3. AHA: AHA added, No AHA: No AHA added, R: Replicate

|  | AHA R1 | AHA R2 | AHA R3 | No AHA R1 | No AHA R2 | No AHA R3 | p value | BenjaminiHochberg Correction | Ratio_group_2/1 | shapiro_Wvalue |
| --- | --- | --- | --- | --- | --- | --- | --- | --- | --- | --- |
| **5 h** | 8.80E+09 | 7.30E+09 | 9.10E+09 | 7.30E+09 | 1.00E+10 | 9.00E+09 | 0.73 | 0.73 | 1.04 | 0.88 |
| **1 h** | 3.80E+09 | 4.70E+09 | 4.70E+09 | 2.15E+09 | 4.60E+09 | 2.70E+09 | 0.23 | 0.46 | 0.72 | 0.83 |

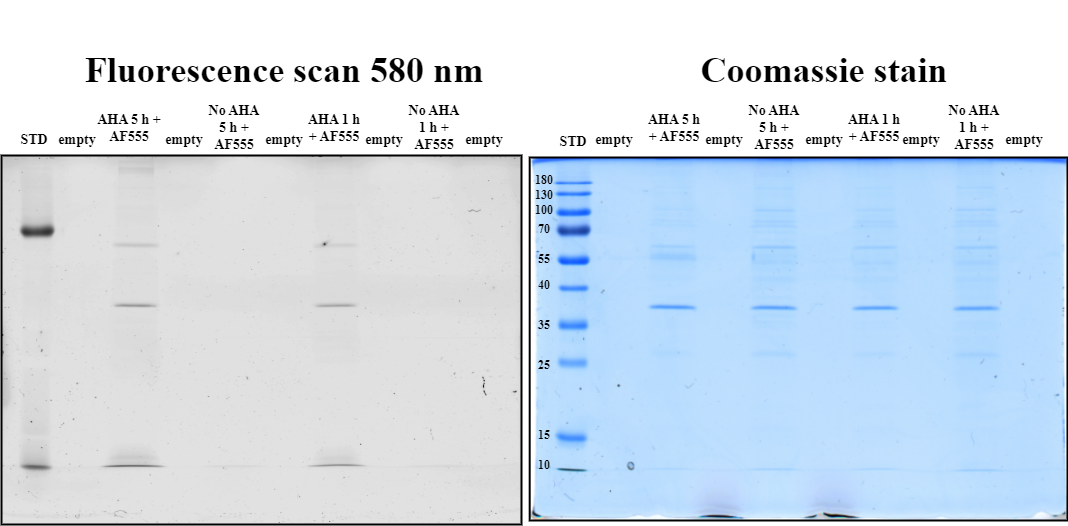

Figure S 1: Fluorescence scan of SDS gel of AHA-labelled phage proteins with DBCO-AF555. The AF555 phages were recovered from 100 kDa filter units. 100 µL of the AF555 phages was precipitated with methanol-chloroform and loaded onto the gel (Material and Methods). AHA: AHA added, 1 or 5 h: incubation time with AHA, AF555: Alexafluor 555, STD: protein standard (Thermo Scientific, PageRuler Prestained Protein Ladder #26616).

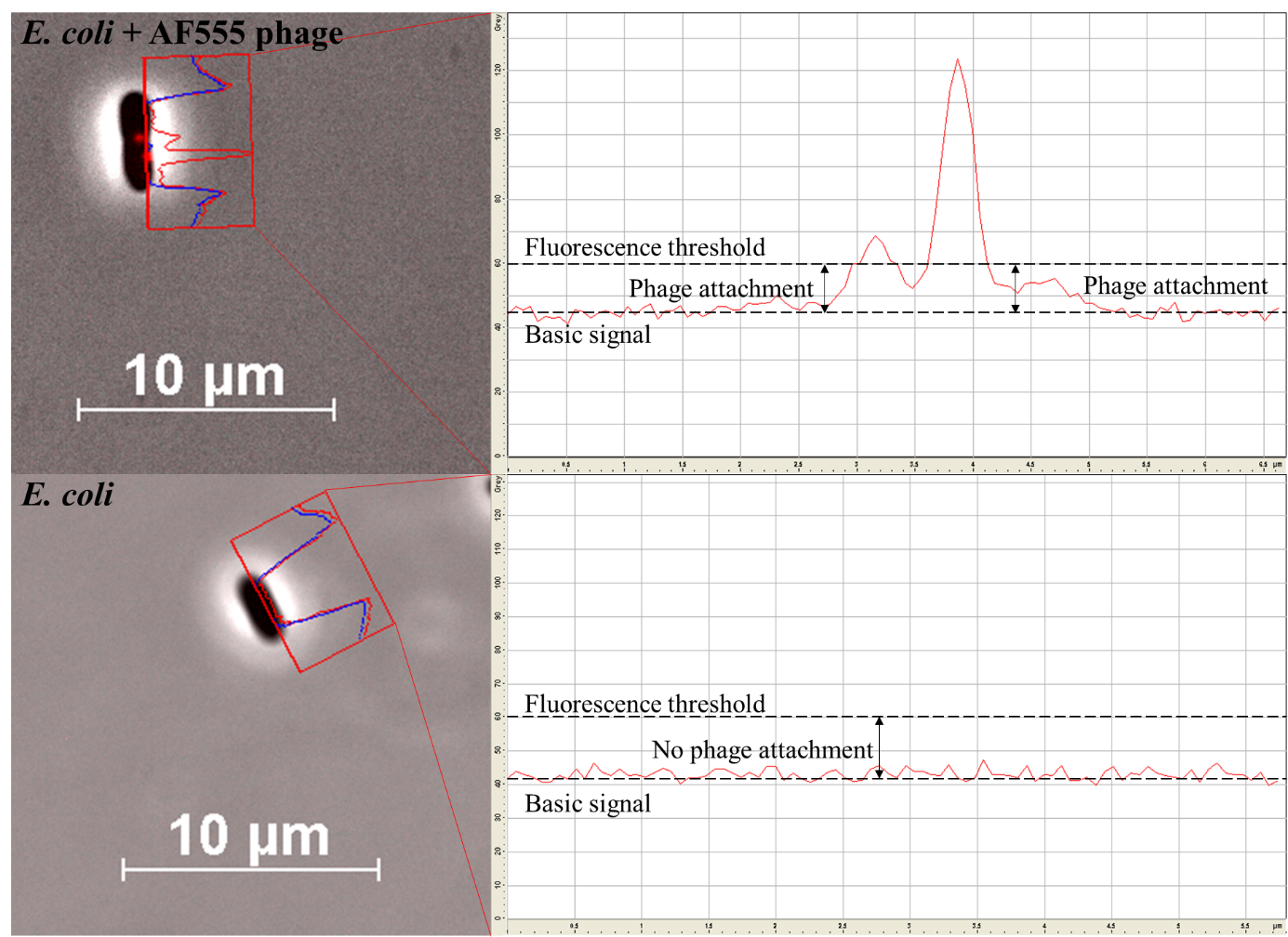

Figure S 2: Counting fluorescence positive phage-host complexes using the greyscale function of the software package AxioVision (Version 4.8.2 SP3). The basic signal of *E. coli* in the fluorescence channel was around 45 arbitrary units (red line). For *P. fluorescens* the basic signal was the same (not shown). The threshold for a positive fluorescence signal was set to 60 arbitrary units. The blue line represents the signal in the brightfield channel.

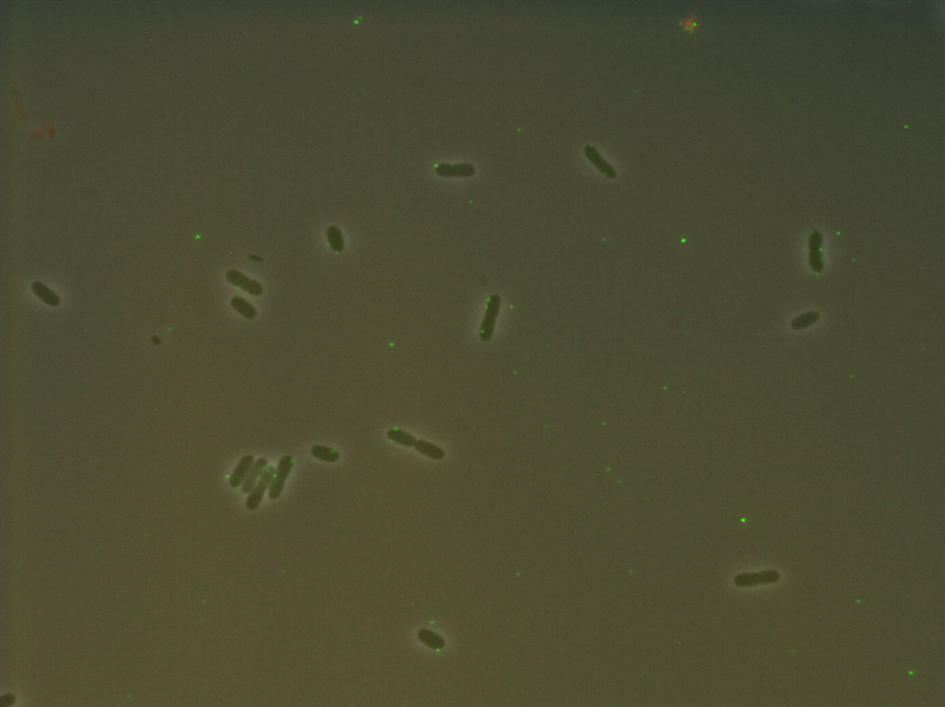

Figure S 3: Fluorescence microscopy of *E. coli* with fluorescent phage λs. The cells were incubated for 30 min with AF488 phage λs (DBCO Alexfluor 488 labelled). The exchange of the fixation solution with PBS was skipped (see Material and Methods). The picture was taken with an Imager.M1 fluorescence microscope (Carl Zeiss, Jena) and the software package AxioVision (Version 4.8.2 SP3). The pictures were taken with brightfield and fluorescence (excitation 470/40 nm; beamsplitter: FT 495; emission 525/50 nm) with a 1000x zoom and phase contrast.

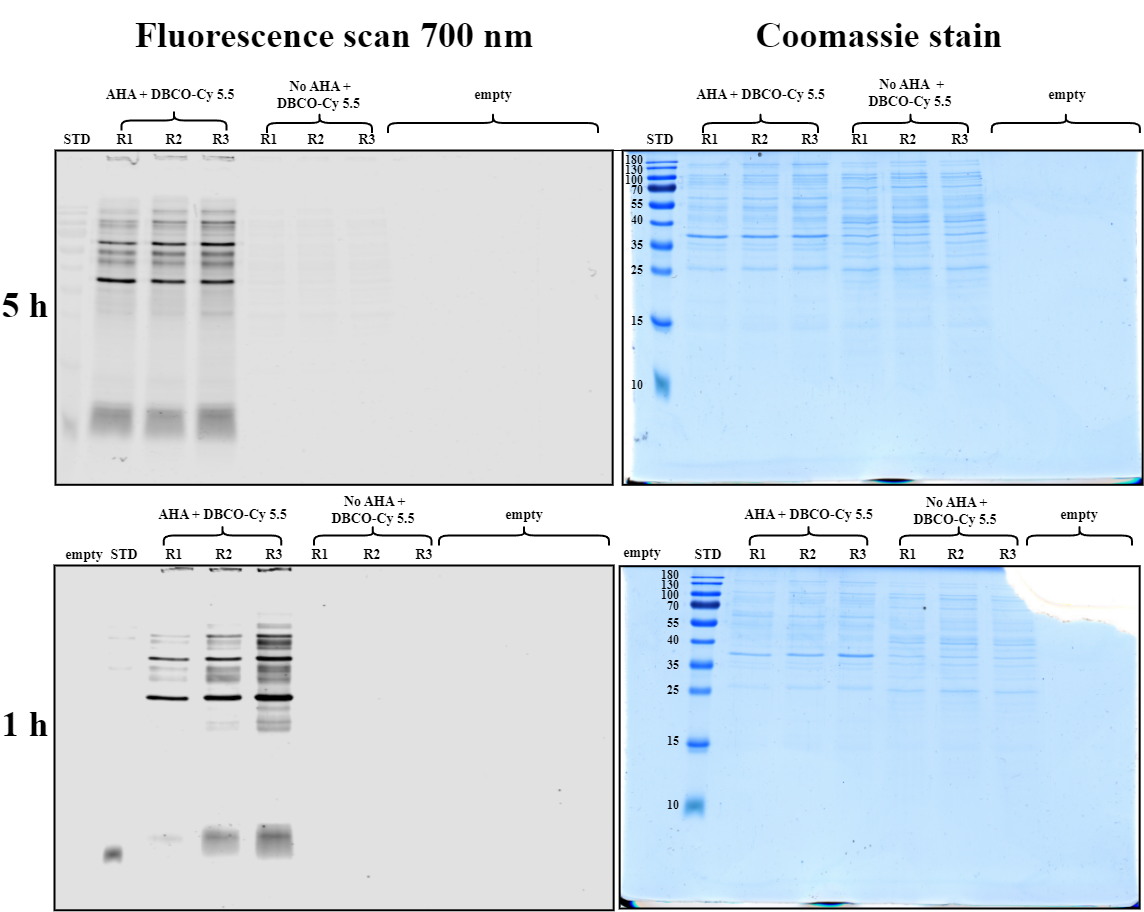

Figure S 4: Fluorescence scan of SDS gel from CY 5.5 phages. AHA: AHA added, STD: protein standard (Thermo Scientific, PageRuler Prestained Protein Ladder #26616)

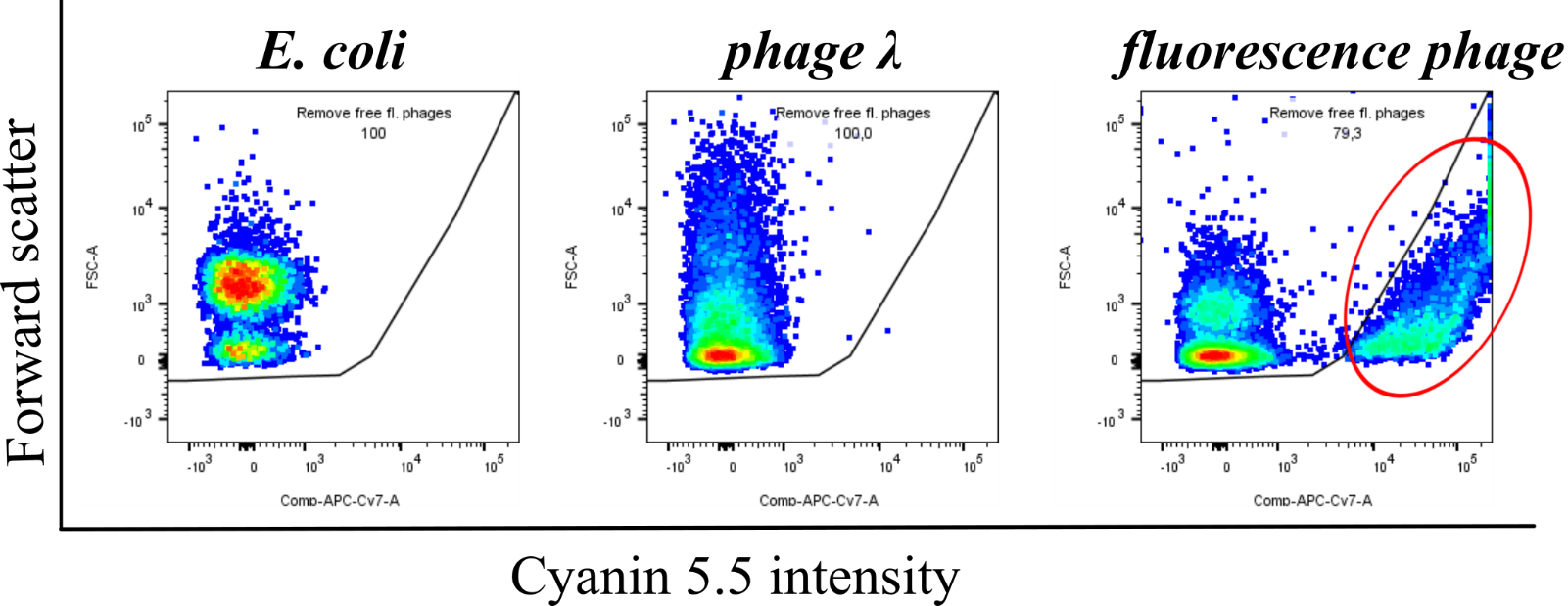

Figure S 5: Gating strategy to remove freely suspended and clumped CY5.5 phages in the background (oval red circle) for discrimination of CY5.5 phages attached to the host cell. The gating was done after analysing the samples with the program FlowJo™ (10.8.1). The gating was adopted for all measurements in Figure 5 (chapter: Quantification of fluorescent phage-host complexes via flow cytometry).

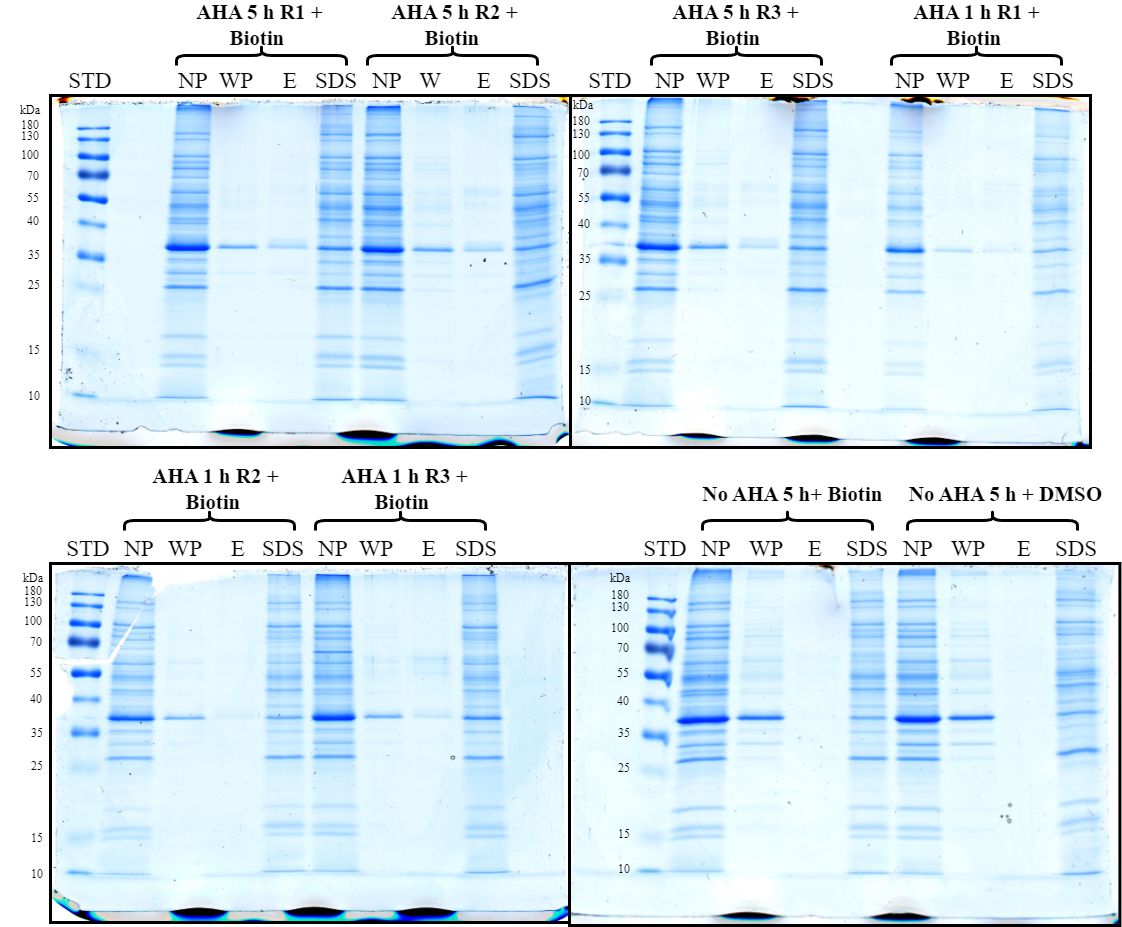

Figure S 6: Purification of biotinylated phages with monomeric avidin beads (all gels). SDS gel of phages incubated with DBCO-PEG4-biotin stained with Coomassie blue. STD: protein standard (Thermo Scientific, PageRuler Prestained Protein Ladder #26616); NP: not purified (no beads added); WP: Wash phase with PBS buffer; E: Elution with 2 mM biotin; SDS: Beads boiled with SDS sample buffer All experiments were performed in triplicates except for the controls with unlabelled phages.
