## Supplementary_note_2_Material_and_Methods_SOPs for "Detection, isolation and characterisation of phage-host complexes using BONCAT and click chemistry"

**Protocol S1 - Induction and production of AHA-labelled phages**

Adapted from Clokie and Kropinski (2009), Hatzenpichler and Orphan (2016) and Pasulka *et al.* (2018)

Solutions:

0.1 M Calcium chloride stock solution:

- 3.78 g CaCl_2_ dihydrate
- Add to 250 mL with water
- Autoclave or filter sterile

1 M Magnesium sulfate stock solution:

- 30.092 g MgSO_4_
- Add to 250 mL with water
- Autoclave or filter sterile

Thiamin stock solution (prepare fresh):

- 100 mg thiamin hydrochloride
- 1 mL pure water
- Filter sterile

M9 minimal medium:

- 0.5 g NaCl
- 1.0 g NH_4_Cl
- 12.8 g Na_2_HPO_4_ × 7 H_2_O
- 3.0 g KH_2_PO_4_
- Add to 1 L with pure water
- Autoclave sterile
- Add 1 mL 1 M MgSO_4_ (sterile)
- Add 1 mL 0.1 M CaCl_2_ (sterile)
- Add 10 mL 10% (w/v) glucose (sterile)
- Directly before cultivation: add 100 mg/L thiamin (end concentration, sterile filtrated)

AHA stock solution (10 mM):

- 1.81 mg 4-Azido-L-homoalanine HCl (Jena Bioscience GmbH, CLK-AA005)
- 1 mL pure water
- Filter sterile

Mitomycin c (MMC) stock solution 1 mg/mL:

- 1 mg MMC (Alfa Aesar, J63193.MA)
- 1 mL Dimethyl sulfoxide (DMSO)

1 M NaOH:

- 2 g Sodium hydroxid (NaOH)
- Add to 50 mL with pure water

Materials:

- 50 mL glass flasks
- Flasks Shaking Incubator for 30-37 °C
- Photometer
- Pipettes (1 -1000 µL)
- 25 mL Serological pipettes
- Centrifuge for 50 ml reaction tubes
- pH Meter
- Syringes with needle
- Syringe filter (5 µm, 1.2 µm, 0.8 µm, 0.45 µM)
- 50 mL reaction tubes (we suggest protein LoBind® Tubes (Eppendorf, Germany) for storage of phages)

Bacteria strains:

- *Escherichia coli* DSM 5911 (<https://www.dsmz.de/collection/catalogue/details/culture/DSM-5911>)
- *Escherichia coli* DSM 5911 with integrated Escherichia phage lambda (*E. coli* plus lambda, kindly provided by Prof. Dr. Andreas Kuhn, University of Hohenheim)

Procedure:

1. Inoculate each 100 mL M9 medium with *E. coli* and *E. coli* plus phage λ in 50 mL flasks
2. Incubate at 37 °C and 130 rpm in a Flasks Shaking Incubator overnight
3. Measure the OD_600nm_ of each culture
4. Use the overnight cultures as an inoculum for the experiment. Prepare a starting OD_600nm_ of 0.1-0.2 for each culture:
   a. Cultures in our experiment per day (perform independent replicates on different days):

i. 7 x 50 mL *E. coli* plus phage λ in M9 medium (OD_600nm_ 0.1-0.2)

ii. 1 x 50 mL *E. coli* (OD_600nm_ 0.1-0.2)

5. Incubate at 37 °C and 130 rpm for 1 h
6. Add 25 µL MMC stock solution (0.5 µg/mL end concentration), 0.5 mL 10 mM AHA (0.1 mM end concentration) and measure the OD_600nm_ over the time as described in Table M1:

Table M 1: Overview of conditions for induction and production of BONCAT phages in *E. coli* K12

| time [h] | A, *E. coli* K12+λ (virus production, AHA 6 h) | B, *E. coli* K12+λ (virus production) | C, *E. coli* K12+λ (AHA 6 h) |
| --- | --- | --- | --- |
| 0 | MMC, AHA, OD_600_ | MMC, OD_600_ | AHA, OD_600_ |
| 1 – 5 | OD_600_ | OD_600_ | OD_600_ |
| 6 | OD_600_, harvest | OD_600_, harvest | OD_600_, harvest |
| time [h] | **D, *E. coli* K12+λ (virus production, AHA 1 h)** | **E, *E. coli* K12+λ (virus production)** | **F, *E. coli* K12+λ (AHA 1 h)** |
| 0 | MMC, OD_600_ | MMC, OD_600_ | OD_600_ |
| 1 | OD_600_ | OD_600_ | OD_600_ |
| 2 | OD_600_ | OD_600_ | OD_600_ |
| 3 | AHA, OD_600_ | OD_600_ | AHA, OD_600_ |
| 4 | OD_600_, harvest | OD_600_, harvest | OD_600_, harvest |
| time [h] | **G, *E. coli* K12+λ (Control strain)** | **H, *E. coli* K12 (Control MMC)** | |
| 0 | OD_600_ | MMC, OD_600_ |  |
| 1 - 5 | OD_600_ | OD_600_ |  |
| 6 | OD_600_, harvest | OD_600_, harvest |  |

7. Harvest of virus: centrifuge the culture at min 3,000 ×g at 4 °C for 12 min.
8. Collect phage containing supernatant and transfer in 50 mL reaction tubes
9. Neutralise the supernatants to pH 7.0 with 1 M NaOH.
10. Filter the supernatants through a membrane filter cascade of 5 µm, 1.2 µm, 0.8 µm, 0.45 µM into a new sterile 50 mL reaction tube
11. Store the sterile supernatant at 4 °C.

**Protocol S2 – Plaque assay**

Adapted from Clokie and Kropinski (2009), Adams (1959), Gratia (1936) and Gratia (2000)

Solutions:

1 M Magnesium sulfate stock solution:

- 30.092 g MgSO_4_
- Add to 250 mL with water
- Autoclave or filter sterile

LB-Medium:

- 10 g NaCl
- 5 g Yeast extract
- 10 g Tryptone
- Add to 1 L with water
- Autoclave

Underlay agar:

- 10 g NaCl
- 5 g Yeast extract
- 10 g Tryptone
- 15 g Agar Agar
- Add to 1 L with pure water
- Autoclave
- Add 5 mL 1 M MgSO_4_ (5 mM end concentration)

Overlay agar:

- 2.5 g NaCl
- 1.25 g Yeast extract
- 2.5 g Tryptone
- 1 g Agar Agar
- Add to 250 mL with pure water
- Autoclave
- Add 1.25 mL 1 M MgSO_4_ (5 mM end concentration)

Materials:

- 50 mL glass flasks
- Flasks Shaking Incubator
- Pipettes (1 -1000 µL)
- Sterile 1.5 mL reaction tubes
- Petri dishes
- Water bath
- Screw-capped glass bottles with at least 4 mL volume

Bacteria strains:

- *Escherichia Coli* DSM 5911 (<https://www.dsmz.de/collection/catalogue/details/culture/DSM-5911>)
- Escherichia phage lambda stock (see Protocol S1)

Procedure:

All steps need to be performed under sterile conditions! Note: Dilution series can be done in a smaller scale, but we suggest the presented volume for higher accuracy.

A. Preparing of Petri dishes with underlay agar

1. Warm underlay agar to 50 °C in a water bath
2. Dispense 25 mL of underlay agar to each Petri dishes
3. Cool the Petri dishes at RT for 30 min and store at 4 °C until use

B. Preparing Overlay agar

1. Aliquot each 4 mL of overlay agar into Screw-capped glass tubes
2. Store at 4 °C (autoclaving before use needed)
3. Warm to 50 °C in a water bath before use

C. Double Agar Overlay Plaque Assay

1. Prepare a culture of *E. coli* in LB-medium
2. Incubate at 37 °C and 130 rpm over night
3. Prepare the overlay agar and hold the 4 mL aliquots in screw-capped glass tubes at 50° C until use
4. Remove the required number of underlay agar plates from 4 °C storage and dry them uncovered in a laminar flow hood for 15 min
5. Label the plates with the phage dilutions to be analysed (here we used 10^-6^-10^-7^per phage stock) and one control (only *E. coli*)
6. Prepare sterile 1.5 mL reaction tubes for the dilution series of the phage stock (here up to 10^-7^, 7 x 1.5 mL reaction tubes per phage stock)
7. Fill 900 µL LB-medium in each 1.5 mL tube
8. Add 100 µL phage stock in the first tube (10^-1^), mix, transfer 100 µL from 10^-1^ into the next tube (10^-2^). Precede until dilution 10^-7^
9. The phage dilution series can be used directly or be stored at 4 °C
10. Take out one flask with overly agar, add 100 µL of the selected phage dilution (e.g. 10^-7^) and 100 µL *E. coli* overnight culture, mix well
11. Pour the overlay agar with the phages and the *E. coli* over the underlay agar in the Petri plate
12. Distribute the liquid overly agar over the plate by swirling gently
13. Repeat for each phage dilution of interest
14. Dry the plates with a partially open lid in the laminar flow hood for 15 min
15. Incubate at 37 °C for 24 h
16. Count plaques on plates, for calculation of phage titer only use plates with 30-300 plaques
17. Number of plaques × 10 × reciprocal of dilution = pfu/mL

**Protocol S3 – Analysis of AHA incorporation into newly produced phage proteins via mass spectrometry**

Adapted from Wessel and Flügge (1984), Heyer *et al.* (2019), Popov (1975), Racusen (1973) and Wiśniewski *et al.* (2009)

Note: The protocols for protein quantification with Amido Black (other protein quantification methods are also possible here) and proteolytic FASP digestion are not described here and can be found in Heyer *et al.* (2019) (<https://www.frontiersin.org/articles/10.3389/fmicb.2019.01883/full#supplementary-material> DATA SHEET S1). Acetone precipitation can be used instead of methanol-chloroform protein extraction. Avoid reducing agents (like DTT, mercaptoethanol, etc.) if a click chemistry reaction is to be carried out afterwards!

Solutions/Chemicals:

0.1 M Tris-HCl, pH 8.5:

- 15.7 g Tris-HCl
- 800 mL pure water
- Adjust pH to 8.5 (with 1 M NaOH (see S1))
- Add to 1 L with pure water

8 M urea buffer:

- 4.8 g urea
- Add to 10 ml with 0.1 M Tris-HCl pH 8.5

Loading A:

- 100 mL LC-MS grade water
- 0.1 mL Trifluoroacetic acid (LC-MS grade)

Methanol

Chloroform

Materials/Chemicals used before LC-MS/MS measurement:

- Centrifuge
- Vacuum centrifuge (Digital Series SpeedVac SPD121P, Thermo Scientific, Waltham, United States)
- 50 mL reaction tubes
- Pipettes with tips (1-10 mL)
- Autosampler Vials and caps for HPLC

A. Protein extraction (perform all steps under a fume hood)

1. Add 4 mL of each phage stock to a new 50 mL reaction tube tube.
2. Add 16 mL methanol to each 50 mL reaction tube tube.
3. Add 4 mL chloroform to each 50 mL reaction tube tube.
4. Add 12 mL pure water to each 50 mL reaction tube tube.
5. Mix well.
6. Centrifuge at 10,000 ×g and room temperature for 5 min.
7. Now, each reaction tube has three phases (lower phase: chloroform, thin middle phase: proteins (!) and upper phase: DNA, lipids, etc.).
8. Carefully remove the upper phase with a pipette.
9. Add 16 mL methanol to each 50 mL reaction tube.
10. Mix well.
11. Centrifuge at > 10,000 ×g and room temperature for 10 min (if higher g is possible, this could help the pellet to be a little firmer).
12. Remove the supernatant by decanting carefully.
13. Let the pellets dry under a fume hood.
14. Add 1 mL 8 M Urea and resuspend the protein pellet.
15. Store at -20 °C till further use.

B. Protein quantification with amido black

- See Heyer *et al.* (2019), <https://www.frontiersin.org/articles/10.3389/fmicb.2019.01883/full#supplementary-material> DATA SHEET S1, Protein quantification with amido black

C. FASP digestion

- 25 µg protein per sample were used
- See Heyer *et al.* (2019), <https://www.frontiersin.org/articles/10.3389/fmicb.2019.01883/full#supplementary-material> DATA SHEET S1, FASP digestion

D. Peptide preparation for LC-MS/MS measurement

1. Dry the peptides generated in C in a vacuum centrifuge.
2. Add 75 µL Loading A to the peptides.
3. Mix well and centrifuge at 2,000 ×g room temperature for 30 sec.
4. Transfer the peptides to HPLC vials and cap the vials.

E. LC-MS/MS measurement

Here, the further procedure depends strongly on the LC-MS/MS system available. In the following, the procedure is described using the LC-MS/MS system specified below as an example.

The peptides were analysed by LC-MS/MS using an UltiMate® 3000 nano splitless reversed-phase nanoHPLC (Thermo Fisher Scientific, Dreieich) coupled online to a timsTOF™ pro mass spectrometer (Bruker Daltonik GmbH, Bremen). The peptides were loaded isocratically on a trap column (Dionex Acclaim, nano trap column, 100 μmi.d. x 2 cm, PepMap100 C18, 5 μm, 100 Å, nanoViper) with a flow rate of 15 μL/min solvent A (100% LC-MS water, 0.1% TFA). For the chromatographic separation, a Dionex Acclaim PepMap C18 RSLC Nano-reversed phase column (2 μm particle size, 100 Å pore size, 75 μm inner diameter, and 500 mm length) with a column temperature of 60 °C was used. First, a 400 nL/min flow rate was applied for a binary solvent A/B gradient (solvent A: 100 % ultrapure water, 0.1 % formic acid; solvent B: 100 % acetonitrile, 0.1% formic acid). Next, 3 µL of the peptide solutions equivalent to ≈1 µg of peptides was injected. The separation step started with 2 % solvent B for 5 min and the concentration of solvent B was linearly increased over a 120 min gradient to 35 %. Afterwards, the column was washed with 95 % solvent B for 10 min and re-equilibrated with 5 % loading B for 20 min. A positive mass spectrometric measurement was combined with a data-dependent MS/MS method. Parallel Accumulation Serial Fragmentation (PASEF) scan mode was performed using trapped ion mobility spectrometry (TIMS) from a range of 0.6 to 1.6 V*s/cm2, charge state from 0 to 5 and 10 PASEF MS/MS scans per cycle. MS/MS data acquisition was performed over the mass range from 100 to 1700 m/z using collision-induced dissociation (CID) with an active exclusion of the same precursor after 1 spectra for 24 seconds or a release if intensity/previous intensity exceeded 4 (Meier *et al.*, 2015). The resulted fragment ions were analysed in a TOF analyser (Mitchell Wells and McLuckey, 2005).

F. Identification of AHA incorporation

The further procedure depends strongly on the data analysis software available for LC-MS/MS data. In the following, the procedure is described using the LC-MS/MS system with Compass DataAnalysis software (version 5.3.0, Bruker Corporation, Bremen, Germany) and MASCOT Daemon (Version 2.6.0) (Perkins et al., 1999) as an example.

1. Raw data files from LC-MS/MS were processed by the Compass DataAnalysis software (version 5.3.0, Bruker Corporation, Bremen, Germany) and converted into Mascot Generic Files (.mgf).
2. UniProt database (.fasta) with only *E. coli* K12 (https://t.ly/4zWd, 23.03.2023) and Escherichia phage lambda (λ phage) (https://t.ly/svCja, 23.03.2023) entities were generated.
3. The .mgf files of each measurement were searched with the following search parameters:

Table M 2: Search parameter for MASCOT MS/MS search

| **Database** | UniProt |
| --- | --- |
| **Taxonomy** | *taxonomy_id: 83333 (23.03.2023) and taxonomy_id: 10710 (23.03.2023)* |
| **Fixed modification** | Carbamidomethylation (M, +57.021464 Da) |
| **Variable modification** | Oxidation (+ 15.994915 Da), L-AHA (M, - 4.986324 Da),  reduced L-AHA (M, - 30.976822 Da) |
| **Enzyme** | Trypsin |
| **Missed Cleavage** | 1 |
| **Peptide charge** | 2+, 3+ and 4+ |
| **Peptide tolerance** | 0.02 Da |
| **MS/MS tolerance** | 0.02 Da |
| **FDR** | <0.01 |

**Protocol S4 – Click chemistry with AHA-labelled phages, phage-host complex detection, and native phage purification.**

Adapted from Hatzenpichler and Orphan (2016), Kolb, Finn and Sharpless (2001) and Pasulka *et al.* (2018)

Notes: The concentration of Iodoacetamide (IAA) can further be optimized. The used fluorescent dyes are examples and were chosen based on the subsequent analysis. Any fluorescent dye can be coupled to AHA as long as it is DBCO-modified for the click reaction. The procedure is described using FACS and fluorescence microscope for detection of labelled phages as an example in the following. Work under sterile conditions if it is needed to store the biotin/dye tagged phages for a longer time (weeks to months). Also, the protocol could be upscaled for bigger filter units or in general, with higher volumes of phage suspensions. Furthermore, a larger pore size (e.g. 300 kDa) of the filters might reduce the background proteome.

Solutions:

1 M HCL:

- 91.66 mL pure water
- 8.33 mL HCl

PBS:

- 8 g NaCl
- 0.2 KCl
- 1.44 g Na_2_HPO_4_
- 0.24 g KH_2_PO_4_
- Add 800 mL pure water
- Adjust pH to 7.4 with 1 M HCl
- Fill up to 1 L with pure water

100 mM Iodoacetamide (IAA) (light-sensitive!):

- 462.4 mg IAA
- 25 mL PBS
- Always prepare fresh!

10 mM Dibenzylcyclooctyne (DBCO) Alexafluor 555 stock solution (light-sensitive!):

- 1 mg DBCO Alexafluor 555 (Jena Bioscience GmbH, CLK-093-1)
- Add 90.5 µL DMSO
- Aliquot and store dark at -20 °C (we use brown 1.5 tubes or use aluminium foil)

10 mM DBCO Cyanin 5.5 stock solution (light-sensitive!):

- 1 mg DBCO Cyanin 5.5 (Jena Bioscience GmbH, CLK-1046-1)
- Add 85.5 µL DMSO
- Aliquot and store dark at -20 °C (we use here brown 1.5 tubes or aluminium foil)

1 M Magnesium sulfate stock solution:

- 30.092 g MgSO_4_
- Add to 250 mL with pure water
- Autoclave or filter sterile

LB-Medium (plus 5 mM MgSO_4_):

- 10 g NaCl
- 5 g Yeast extract
- 10 g Tryptone
- Add to 1 L with water
- Autoclave
- Optional: Add 5 mL 1 M MgSO_4_ for possibly better phage adsorption

Fixation solution:

- 4.559 mL PBS
- 0.541 mL 37 % formaldehyde solution
- Prepare always fresh!

10 mM DBCO-PEG4-Biotin stock solution:

- 10 mg DBCO-PEG4-Biotin (Jena Bioscience GmbH, CLK-A105P4-10)
- 1.334 mL DMSO
- Aliquot and store dark at -20 °C

Regeneration Buffer (0.1 M Glycine/HCl, pH 2.8):

- 150.14 mg Glycine
- 15 mL with pure water
- Adjust pH to 2.8 with 1 M HCl
- Fill up to 20 mL with pure water
- Store at 4 °C till further use

Blocking/Elution Buffer (2 mM D-biotin in PBS):

- 9.77 mg D-biotin
- 20 mL PBS
- Store at 4 °C till further use

Materials:

- 4 mL 100 kDa filter unit (Amicon Ultra-4, PLHK Ultracel-PL Membrane, 100 kDa)
- Centrifuge for Reaction tubes
- pH meter
- Pipette with tips (1-1000 µL)
- Serological pipettes
- Sterile 1.5 mL tubes (we suggest LoBind® Tubes (Eppendorf, Germany) for storage of phages)
- Photometer (ThermoScientific, Genesys 10S UV-VIS)
- Thermal shaker (for 1.5 mL tubes)
- Warming cabinet (Incubator) for 30-40 °C
- BcMag™ Monomeric Avidin Magnetic beads (Bioclone Inc, MMI-102)
- Magnetic rack (e.g., https://www.jenabioscience.com/molecular-biology/rna-dna-preparation/magnetic-racks/pp-229-magrack-6)

Test strains:

- *Pseudomonas fluorescens* (DSM 50090) → control strain, cannot be infected by lambda phages
- *Escherichia coli* (DSM 5911) → host of lambda phages

Procedure:

A. Click chemistry with AHA labelled phages (from protocol S1)

1. Add aimed phages to the 100 kDa filter unit (we used 4 mL per phage stock).
2. Centrifuge at 3,500 ×g and RT (or longer if necessary) for 5 min.

IMPORTANT: Let always at least 50-100 µL liquid on the filter; otherwise, high losses in phage titer can occur

3. Wash 1x with 4 mL PBS and centrifuge for 5 min, 3,500 ×g, RT (or longer if necessary).
4. Add 4 mL 100 mM IAA in PBS per sample.
5. Mix well.
6. Incubate 1 h at 37 °C in the dark with light shaking.
7. Dilute stock solution of fluorescent dye (e.g. 10 mM DBCO Cyanin 5.5) with DMSO or water to 1 mM, e.g. 1 µL stock + 9 µL DMSO, the DBCO-PEG4-Biotin stock solution does not need to be diluted.
8. Add 0.6 µL 1 mM fluorescent dye or 60 µL 10 mM DBCO-PEG4-Biotin stock solution to the filters.
9. Mix well.
10. Incubate at 37 °C in the dark with light shaking (up to 1 h is possible) for 30 min; the reaction can also be done overnight at 4 °C.
11. Optional: Add 1 mM AHA (end concentration) to the phages and incubate for 10 min at 37°C to stop the reaction (we skipped this step because we immediately continued with centrifugation).
12. Centrifuge at 3,500×g and RT for 5 min (or longer if necessary).
13. Add 4 mL PBS.
14. Mix well.
15. Centrifuge for 5 min, 3,500×g, RT (or longer if necessary)
16. Repeat step 13.-15. three times.
17. Add 1 mL PBS.
18. Mix well, as phages tend to stick to the filter; it is crucial to detach them.
19. Transfer the phage suspension into a sterile 1.5 mL tube (we suggest LoBind® Tubes (Eppendorf, Germany) for storage phages).
20. Store the tagged phages at 4 °C till use.

B. Optional: Quality control for recovery efficiency, Plaque-assay

- see protocol S2

C. Optional: Quality control for fluorescence, SDS-PAGE with in-gel fluorescence scan

1. Use 100 µL from each phage suspension.
2. Precipitate with ice-cold acetone (100%) at -20 °C for at least 1 hour.
3. Do SDS-PAGE, See Heyer *et al.* (2019), <https://www.frontiersin.org/articles/10.3389/fmicb.2019.01883/full#supplementary-material> DATA SHEET S1, SDS-PAGE.
4. Fixate the gels with fixation solution for 1 h at RT in the dark
5. After fixation of the gel, scan the fluorescence of the proteins within the gel (for settings, see table below).
6. Stain the gel with Coomassie following the manufacturer´s description.
7. Scan gel with a gel scanner (for settings, see table below).

Table M 3: Parameter for scanning SDS-gels and fluorescence

| Stain/Fluorescence | Scanner | Software | PArameter |
| --- | --- | --- | --- |
| DBCO-Cyanin 5.5 | Licor Odyssey ODY-2600 (LI-COR Biosciences - GmbH) | Image Studio™ (Version 5.2.5) | 700 nm Wavelength, intensity: 3, resolution 169 µm, quality: medium, analysis: manual |
| DBCO-Alexafluor 555 | Typhoon Trio Variable Mode Imager System (GE Healthcare) | Typhoon Scanner Control (Version 5.0) | Laser 532 nm and emission filter 580 nm, 600 V, Acquisition Mode: Fluorescence, Pixel size: 100 microns, Focal Plane: Platen |
| Comassie | Biostep ViewPix900 scanner (Seiko Epson Corporation) | ArgusX1 (Version 7.4.28) | Scan mode: Transmission  Colour depth: 48-bit  Brightness: 0  Contrast: 0  Gamma: 1.2  red: 100, green: 100 and blue:100 |

D1. Adsorption of fluorescence-tagged phages to host cells:

1. Prepare 50 mL overnight culture in LB-medium (plus 5 mM MgSO_4_) from *P. fluorescens* and *E. coli.*
2. Incubate at 130 rpm and 30 °C overnight (temperature where both strains can grow)
3. Measure OD_600nm_ with a photometer.
4. If a specific Moi is needed later, a program can be used to get an idea of the *E. coli*/*P. fluorescence* concentration: <https://www.agilent.com/store/biocalculators/calcODBacterial.jsp>. However, this approach is not possible for all bacteria strains and not for mixed cultures. For other strains, flow cytometric analysis (Bailey *et al.*, 1977; Paau, Cowles and Oro, 1977) or koch's plate pour method (Gerhardt, 1994) are possibilities to determine the cell number. For unknown cultures different dilutions of a cell sample should be used for the later phage adsorption.
5. *Dilute E. coli* and *P. fluorescens* to an OD_600_ of 0.02 (≈ 1.80 × 10^7^ cells/mL).
6. Prepare and label 1.5 mL tubes for the adsorption experiment (here per replicate: 2x pure bacteria (*P. fluorescens* and *E. coli*), 2x *E. coli* + lambda phage (1x incubated with fluorescent phages and 1x incubated without fluorescent phages), 2x *P. fluorescens* + lambda phage (1x incubated with fluorescent phages and 1x incubated without fluorescent phages).
7. Add 1mL of *E. coli* or *P. fluorescens* (OD_600nm_ of 0.02) to the related tubes.
8. Incubate at 30 °C and 600 rpm for 20 min.
9. Add fluorescent or unlabelled phages to the related tubes (here, we aimed a moi of ≈ 2).
10. Add PBS or media to the controls.
11. Incubate at 30 °C and 600 rpm in the dark.
12. Take 200 µl samples at defined time intervals, e.g. 0 min (directly after the addition of phages), 10 min, 20 min, 30 min and 60 min.
13. After each sampling, centrifuge the taken sample immediately at 16,400 ×g and 4 °C for 5 min.
14. Remove carefully the supernatant with a pipette.
15. Add the same volume (as supernatant) of fixation solution (optional).
16. Incubate at 4 °C in the dark for 1 h.
17. Centrifuge at 16,400 ×g and 4 °C for 5 min.
18. Carefully remove the supernatant with a pipette.
19. Add 200 µL PBS.
20. Store phage-hosts complexes at 4 °C. Analyse samples within 3 days.

E1. Fluorescence microscopy

Here, further procedures depend strongly on the fluorescence microscopy setup. In the following, the procedure is described using the fluorescence microscope specified below as an example:

The samples were analysed with an Imager.M1 fluorescence microscope (Carl Zeiss, Jena, Germany) using a 100X objective (EC-Neoflur 100x/1.3 Oil Ph3) and phase contrast. The digital pictures were taken with AxioVision (Version 4.8.2 SP3) and the camera Axiocam MRm (Carl Zeiss, Jena). FITC filter set (excitation 546/12 nm; beamsplitter: FT 560; emission 575-640 nm) with an exposure time of 500 ms was used for fluorescence. For bright field, 110 ms exposure time was used. The cells were scanned manually for fluorescence via the greyscale function of the fluorescence channel. The noise of background and nonfluorescent cells was around 45 on the greyscale. Therefore, the threshold for positive fluorescence was set to 60 on the greyscale (see Figure S 2).

F1. Flow cytometer

Here, the procedure depends strongly on the flow cytometer/FACS setup used. In the following, the procedure is described using the flow cytometer specified below as an example.

Flow cytometric analysis was performed using a FACS Canto II equipped with 3 lasers (405 nm, 488 nm, 663 nm), Firmware Version 1.47 (BD Biosciences, Franklin Lakes, NJ, USA). The flow rate was set to low. *E. coli* and *P. fluorescens* were gated from an FSC-A against an SSC-A dot plot. The binding of fluorescent phages to *E. coli* and *P. fluorescens* was detected in the FITC channel. Fluorescence of *E. coli* and *P. fluorescens* after binding of unlabelled phages and autofluorescence of uninfected *E. coli* served as negative controls. For each experimental condition, 10.000 cells were analysed. Data measurement was performed with FACS Diva, Version 6.1.3. (BD Biosciences). Data were exported as .fcs files and analysed using Flow Jo, Version 10 (BD Biosciences). Pseudocolor plots were used to display the relative population density of cell populations by colours (blue and green for low cell density, yellow for mid-range cell density and red and orange for high cell density).

D2. Purification of biotinylated phages

Adapted manufacturer’s protocol (http://www.bioclone.us/Monomeric-avidin-magnetic-beads-particle-resin-matrix.html)

1. Gently shake the bottle containing BcMagTM Monomer Avidin Magnetic Beads until the magnetic beads are completely suspended. Transfer 100 μl beads per reaction to 1.5 ml reaction tubes.
2. Place the tubes on a magnetic separator for 1 minute. Remove the supernatant while the tubes remain on the separator.
3. Remove the tube from the separator. Add 4 bead volumes of pure water, mix well and place the tube on the magnetic separator for 1 min. Remove the supernatant while the tubes remain on the separator.
4. Add 4 bead volumes of PBS, mix well and place the tubes on a magnetic separator for 1 min. Remove the supernatant while the tubes remain on the separator.
5. Add 3 bead volumes of 1x Blocking / Elution Buffer, mix well by vortexing and incubate at room temperature for 5 min. Place the tube on the magnetic separator for 1 min. Remove the supernatant while the tube remains on the separator.
6. Add 6 bead volumes of 1x Regeneration Buffer, mix well by vortexing and place the tube on the magnetic separator for 1 min. Remove the supernatant while the tube remains on the separator.
7. Add 4 bead volumes of PBS, mix well and place the tubes on a magnetic separator for 1 min. Remove the supernatant while the tubes remain on the separator.
8. The beads are ready to use (Note: The beads must be used immediately, or the binding capacity will be dramatically reduced.)
9. Add 200 µL biotinylated phage suspension, mix well by pipetting and incubate at room temperature with gentle rotation for 60 min.
10. Place the tube on a magnetic separator and remove the supernatant while the tube remains on the separator. Optional: collect supernatant for wash efficiency control, fraction “washing phase”
11. Wash the beads with 1x PBS as in step 2 until the absorbance of the eluate at 280 nm approaches the background level (OD_280 nm_ < 0.05). Optional: collect supernatant for wash efficiency control, combine with supernatant from step 10, fraction “washing phase”
12. Add 200 µL of Blocking/Elution Buffer, mix well by pipetting several times, and incubate at room temperature for 10 min to elute the bound biotinylated phages from the magnetic beads.
13. Place the tubes on a magnetic separator for 1 min. Transfer the supernatant while the tubes remain on the separator. The phages are in the supernatant. Fraction “elution”
14. Optional: Quality control for elution efficiency:
    a. Add SDS sample buffer (see Section C. SDS-PAGE) to the beads and boil the beads for at 1400 rpm and 60 °C 5 min to elute all remaining proteins from the beads. Note that this will destroy the beads! Fraction “SDS-boiled”

E2. Optional: Quality control for elution from D2 step 13, Plaque-assay

- See protocol S2; note dilution series can also be done in small scale, e.g. 20 µL labelled phage + 180 µl media instead of 100 µL phage + 900 µL media.
- We used 100 µL phages.

F2. Optional: Quality control for purification, SDS-PAGE

- Use 100 µL from each biotinylated phage. Additionally, we extracted the proteins from the fraction “washing phase” (from D2 step 10 and 11)
- Use methanol-chloroform precipitation (see protocol S3)
- Fraction “SDS-boiled” (from D2 step 14) were also used for the SDS-PAGE. see the magnetic separator to remove the beads from the supernatant before SDS PAGE (The proteins are in the supernatant)
- Do SDS-PAGE, See Heyer *et al.* (2019), <https://www.frontiersin.org/articles/10.3389/fmicb.2019.01883/full#supplementary-material> DATA SHEET S1, SDS-PAGE
- Fixate the gel for 1 h with fixation solution in the dark.
- Stain with Coomassie overnight.
- Scan gel with a gel scanner (see Table 3).

G. Further analysis of the phages

This strongly depends on the research question and the purpose of using this protocol. Examples of further analysis are sequencing of unknown phages or LC-MS/MS measurement of the phage proteome. We did, as a last step, an in-gel digestion of eluted phages in section F2. to verify the elution of the phages:

- In-gel, See Heyer *et al.* (2019), <https://www.frontiersin.org/articles/10.3389/fmicb.2019.01883/full#supplementary-material> DATA SHEET S1, SDS-PAGE
